# Cortical and white matter myelination proceed in concert during early infancy

**DOI:** 10.1101/2025.07.07.663449

**Authors:** Stephanie Zika, Kelly Chang, Altan Orhon, John Kruper, Christina Tyagi, Xiaoqian Yan, Sarah Tung, Kalanit Grill-Spector, Ariel Rokem, Mareike Grotheer

## Abstract

The infant brain undergoes rapid myelination that is critical for healthy brain function. This development has been characterized for gray and white matter independently, but the link between gray and white matter myelination remains unexplored. To close this knowledge gap, we evaluated two complementary myelin-sensitive imaging metrics: Large-scale (N=273) T1w/T2w and quantitative (N=21) R1 data. Automated software was employed to identify 26 white matter bundles and map their cortical terminations, before evaluating T1w/T2w and R1 development shortly after birth. Here we show that for both metrics mean values as well as developmental slopes are correlated across tissues. The synchrony of brain T1w/T2w is impacted by gestational age and prematurity, whereas inter-individual differences in this synchrony predict motor outcomes at 17 - 25 months of age. As T1w/T2w and R1 are associated with myelin content, our results reveal an intricate relationship between gray and white matter myelination.

## Introduction

The human brain shows rapid development during early life. At birth, total brain volume is about 35% of adult size, growing to nearly 80% by the age of two years old^1^ (for review see ^2^). This remarkable expansion is accompanied by changes in tissue microstructure and organization, including growth of myelin in both gray and white matter^3–6^ (for reviews see ^7–9^). Myelination is a critical process in brain development that involves wrapping neuronal axons in a fatty sheath, thereby enabling rapid and synchronized neural communication. This mechanism is essential for brain plasticity and learning, and disruptions in myelin development have been linked to various developmental and cognitive disorders^10,11^ (for reviews see^12–15^).

In the white matter, myelin insulates axons for rapid saltatory conduction, reduces the energetic cost of signaling by lowering membrane capacitance, supplies metabolic substrates, and preserves axonal structural integrity^16–18^ (for reviews see^19,13^). Although the majority of myelin is located in the white matter^20,21^, a considerable number of myelinated axons can also be found in the gray matter^22,23^ (for review see^24–27^). The precise role of cortical myelin remains debated, with several functions proposed, including i) metabolic support^28^, ii) network synchronization^29,30^ (for review see^31^), iii) fine-tuning of conduction speed^32^, and iv) growth inhibition to prevent aberrant axonal sprouting and synapse formation^33–36^ (for review see^37^).

In both white and gray matter, myelination during early infancy follows a complex spatio-temporal trajectory and, critically, similar mechanisms have been proposed to describe these developmental trajectories across tissues: 1) Spatial gradients: In white matter, several spatial gradients of myelination have been proposed including decreasing developmental rates from: central-to-peripheral^38–41^, posterior-to-anterior^38,5^, and superior-to-inferior^5,42–45^ white matter locations. Similarly, in gray matter, studies revealed a systematic decrease in myelination toward parietal, temporal, and prefrontal cortices^4,22,46–50^. 2) Functional systems: Sensory and motor white matter pathways myelinate earlier than pathways associated with higher-level brain functions^38,41^ (for review see^51^). Similarly, sensory and motor gray matter regions myelinate early and heavy, while higher-order areas such as the prefrontal and association cortices, myelinate later and remain more lightly myelinated even in adulthood^3,4,22,52–57^ (for review see^58^). 3) Myelin content at birth: Recent neuroimaging work suggests that white matter that is less myelinated at birth develops more rapidly postnatally^3,59^ (for review see^60^). Similarly, primary sensory and motor cortices, which exhibit higher myelin levels at term-equivalent age, myelinate more slowly than higher-order association and visual areas during early infancy^4,57,61^ (for review see^62^). Importantly, these mechanisms are likely not mutually-exclusive but rather shape myelin development in concert. Moreover, as the mechanisms that control the spatiotemporal trajectory of myelination in gray and white matter show substantial overlap, there may be an as of yet unexplored link between gray and white matter myelination during infancy.

To test this link, it is critical to follow and compare the trajectories of myelin growth of gray and white matter in the living human brain. Recent advances in quantitative MRI^3,43,63–65^ (qMRI, for review see^66^), have made it feasible to quantify myelin levels and to compare them between individuals and across developmental timepoints. The validity of using qMRI measures such as the myelin-water-fraction (MWF) or the longitudinal relaxation rate (R1), to study myelination has been confirmed by comparisons to post-mortem histological data^67–69^. However, qMRI measures often require long acquisition times, and such measures are hence typically not included in large-scale open data. As a practical alternative, Glasser and Van Essen^22^ proposed utilizing the ratio of T1-weighted (T1w) to T2-weighted (T2w) images to approximate myelin levels. Although the validity of T1w/T2w as a marker of myelin remains debated^70–73^, studies comparing T1w/T2w to R1 have validated its usability in adult cortex^74^ and infant white matter^59^. As the T1w/T2w ratio is easily computed from commonly acquired images, it is particularly useful in large-scale data sets and/or early life samples where acquisition times are particularly restricted.

In this work we combine these approaches to gain both precision and robustness by evaluating a small-scale (N=21) R1 data set collected locally at Stanford University (Stanford VPNL Baby Project (SVBP) data) as well as a large-scale (N=273) T1w/T2w data set shared openly by the Developing Human Connectome Project (dHCP). In both data sets, we test for a link between gray and white matter myelination in early infancy by relating R1 or T1w/T2w of white matter bundles to R1 or T1w/T2w measured at their corresponding cortical targets. We find that white matter and cortical myelin contents are tightly coupled: For both R1 and T1w/T2w values within each bundle are positively correlated with values at its corresponding cortical targets. Similarly, for both R1 and T1w/T2w, the rate of change during early infancy is coupled between white and gray matter. We also observe large inter-individual variability in the strengths of white and gray matter coupling, which, in the dHCP data, is related to gestational age, and correlates with motor performance later in life. In the dHCP data, the coupling between T1w/T2w of white and gray matter is weaker in preterm than full-term infants, even at term-aquivalent age. By highlighting the tight coupling of gray and white matter myelin growths, our findings provide a comprehensive understanding of early life brain myelination and suggest that white and gray matter myelination should be considered conjointly.

## Results

### Coupling of bundles and their cortical targets

We used two complementary myelin-sensitivity imaging metrics - T1w/T2w and R1 - to link development of white matter bundles and their corresponding gray matter targets during early infancy. For this, we leveraged two datasets: i) large-scale data from the Developing Human Connectome Project (dHCP), which contains dMRI and T1w/T2w data from 273 infants, including both preterm and full-term infants, scanned shortly after birth (gestational age (GA) at birth: mean ± SD: 38.05 ± 3.67 weeks; time between birth and scan: mean ± SD: 1.47 ± 2.02 weeks) and ii) locally-collected quantitative data (the Stanford VPNL Baby Project (SVBP) data), which contained dMRI and R1 data from 21 infants scanned shortly after birth (GA at birth: mean ± SD: 39.09 ± 1.63 weeks; time between birth and scan: mean ± SD: 4.30 ± 1.35). In both datasets, we used pyBabyAFQ to identify 26 white matter bundles in individual infants’ brains (Supplementary Figures 1-2), mapped their endpoints to cortex (Fig. 1), and then assessed the development of myelin-sensitive imaging metrics across tissues.

**Figure 1.**
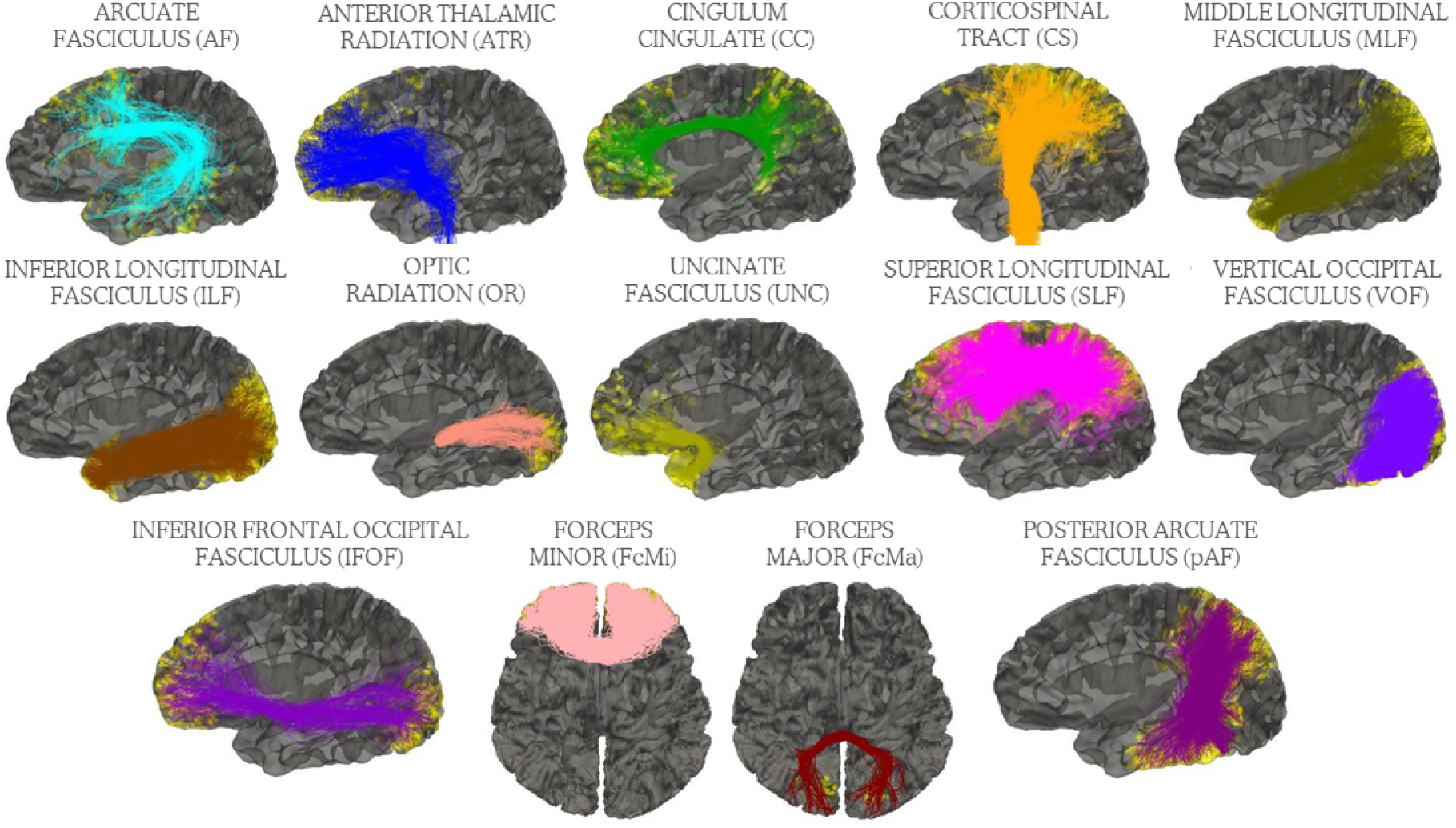
The bundles and their cortical endpoints. Bundles were identified with pyBabyAFQ in the native brain space of one example full-term infant from the dHCP data scanned at 40 weeks GA. The yellow dots represent the cortical endpoints of the respective bundles. For bilateral bundles only the left hemisphere is shown. GA=gestational age.

To investigate the relationship between white and gray matter myelin levels in early infancy, we correlated the myelin-sensitive imaging metrics of white matter bundles with their gray matter targets. In the dHCP data, our analyses of T1w/T2w revealed a positive correlation (Fig. 2a, r^2^ = 0.55, p-value = 1.45e^-5^), indicating that if a white matter bundle has high T1w/T2w, its gray matter targets also have high T1w/T2w. Similarly, in the SVBP data, our analyses of R1 showed a positive correlation (Fig. 2b, r^2^ = 0.32, p-value = 0.002) indicating that if a white matter bundle has high R1, its gray matter targets also have high R1. For both metrics, these observed correlations for the true bundle–target pairings were above the 95% confidence interval (CI) of chance-level correlations (95% CI of chance-level: r^2^=0.14 for T1w/T2w and r^2^=0.12 for R1) obtained by 1,000 iterations of correlating shuffled bundle-target pairings, omitting the true pairs (Supplementary Figure S3).

**Figure 2.**
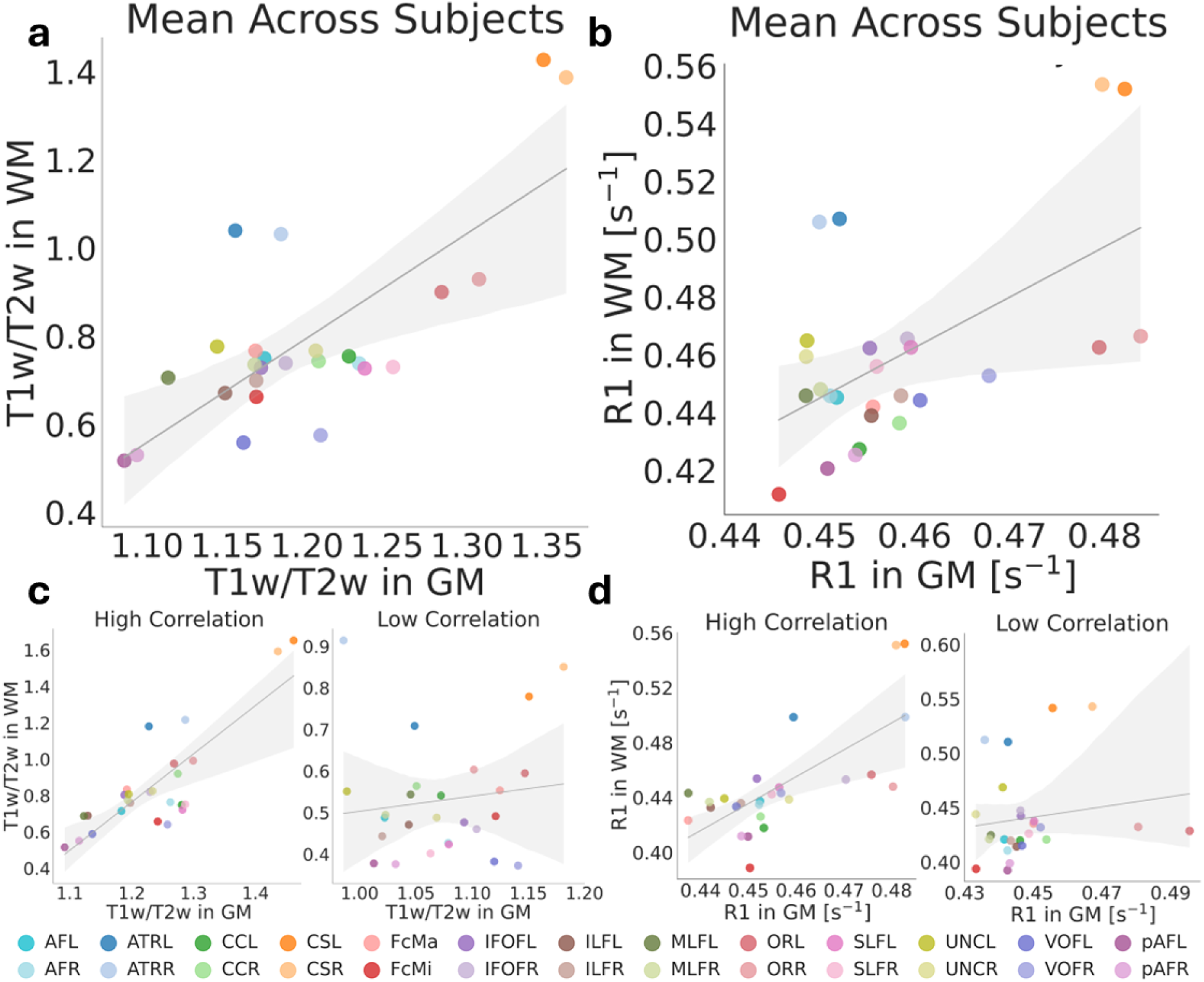
Relationship between T1w/T2w and R1 of WM bundles and their respective GM targets. Each dot is a bundle. **a** Mean correlation with T1w/T2w values averaged across subjects (r^2^=0.55, p-value=1.45e^-5^). **b** Mean correlation with R1 values averaged across subjects (r^2^=0.32, p-value=0.002). **c** Example individual subjects exhibiting high (left; GA at scan: 41.43 weeks; r^2^=0.68, p-value=2.28e^-7^) and low (right; GA at scan: 31.43 weeks; r^2^=0.017, p-value=0.52) correlation between T1w/T2w of white and gray matter. **d** Example individual subjects exhibiting high (left; GA at scan: 42.5 weeks; r^2^=0.51, p-value=4.21e^-5^) and low (right; GA at scan: 40.4 weeks; r^2^=0.03, p-value=0.44) correlation between R1 of white and gray matter. Abbreviations: WM: white matter, GM: gray matter, AF: Arcuate Fasciculus, ATR: Anterior Thalamic Radiation, CC: Cingulum Cingulate, CS: Cortico-Spinal Tract, FcMa: Forceps Major, FcMi: Forceps Minor, IFOF: Inferior Frontal Occipital Fasciculus, ILF: Inferior Longitudinal Fasciculus, MLF: Middle Longitudinal Fasciculus, OR: Optic Radiation, SLF: Superior Longitudinal Fasciculus, UNC: Uncinate Fasciculus, VOF: Ventral Occipital Fasciculus, pAF: Posterior Arcuate Fasciculus, L=left, R=right.

Interestingly, R1 and T1w/T2w were correlated with each other in each tissue across the datasets (Supplementary Figure S4; Gray matter: r^2^=0.66, p-value=5.32e^-7^; White matter: r^2^=0.85, p-value=1.88e^-11^) and revealed similar patterns among different bundles: The corticospinal tract (CS) along with its cortical targets, showed particularly high T1w/T2w and R1. In contrast, the posterior arcuate fasciculus (pAF) and its cortical targets exhibited comparatively low values for both metrics, with other bundles showing intermediate values of T1w/T2w and R1. In the anterior thalamic radiation (ATR) the link between gray and white matter was less pronounced than in the other bundles, with the cortical targets of the ATR having lower values than expected based on their white matter values. Interestingly, our analyses of T1w/T2w and R1 also revealed similar values across hemispheres in gray and white matter of bilateral bundles as indicated by the close proximity of data points representing corresponding bundles in each hemisphere. A detailed overview of differences in T1w/T2w and R1 values across bundles for each tissue is presented in Supplementary Figure S4.

In addition to evaluating the relationship of the mean T1w/T2w and R1 values of gray and white matter at the group-level, we also evaluated this relationship within each individual participant. These individual subjects analyses corroborated the group-level findings, as white and gray matter values were correlated within most individuals for both T1w/T2w (correlation significant in 265 out of 273 subjects (97%), mean r^2^ across subjects: 0.42, mean p-value across subjects: p=0.008) and R1 (correlation significant in 17 out of 21 subjects (81%), mean r^2^ across subjects: 0.26, mean p-value across subjects: p=0.04). Nonetheless, there were large inter-individual differences in the degree of white and gray matter coupling in both T1w/T2w (range of r^2^: 0.01-0.68, standard deviation of r^2^: 0.13) and R1 (range of r^2^: 0.03-0.51, standard deviation of r^2^: 0.12). Figure 2c and 2d show two example individuals each for high and low T1w/T2w and R1 correlations across tissues. Similar to the group level data, in both T1w/T2w and R1, the individual subject analyses showed particularly high gray and white matter values for the CS and comparatively low values for the pAF, with other bundles falling in-between. Further, again similar to the group-level data, the cortical targets of the ATR showed lower T1w/T2w and R1 values than expected based on their white matter T1w/T2w and R1. Finally, in both metrics, bilateral bundles showed similar values in gray and white matter across hemispheres in the individual subject analyses as well.

### Coupled development of bundles and their cortical targets

We further investigated the relationship between myelin-sensitive imaging metrics of white matter bundles and their cortical targets by evaluating the slopes of T1w/T2w and R1 growth during early infancy (Fig. 3). The slopes represent the rate at which each metric changes with infants’ age at scan in the respective age ranges covered by each dataset. Figure 3a–b show example slopes for the corticospinal tract (CS, top) and the cingulum cingulate (CC, bottom); visualizations of all bundles are provided in Supplementary Figure S6 and S7.

**Figure 3:**
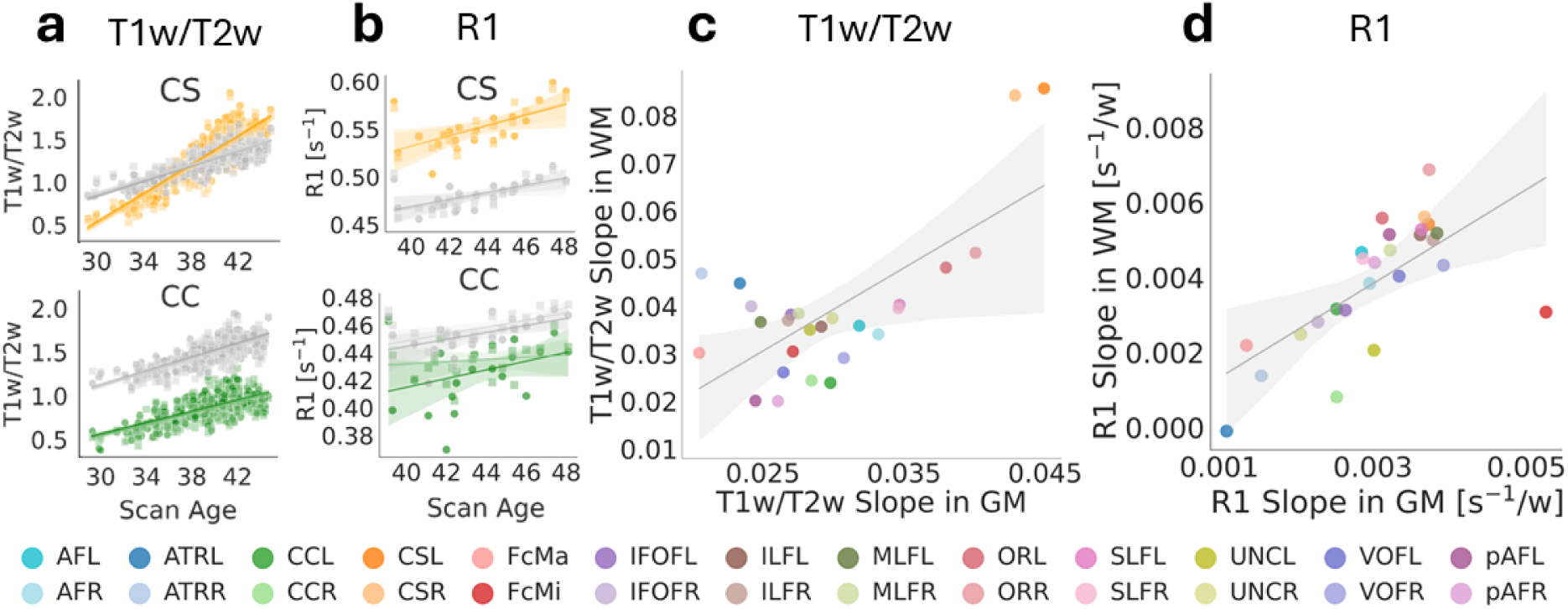
The rates of change of myelin-sensitive imaging metrics are correlated across white and gray matter. a,b. Two example bundles (cortico-spinal tract (CS, top) and cingulum cingulate (CC, bottom)) illustrate T1w/T2w (a) and R1 (b) slopes in gray and white matter (for all bundles see Supplementary Figures S6 and S7). Gray circles indicate gray matter values in each subject, colored circles represent white matter values in each subject, and the steepness of the lines indicate the slopes (rates of change relative to the infant’s gestational age at the time of measurement in weeks), the two hemispheres are presented in different shades. **c,d** Relationship between the slopes of T1w/t2w (c) and R1 (d) along white matter (WM) bundles and their gray matter (GM) targets (T1w/T2w: r^2^=0.49, p-value=6.28e^-5^; R1: r^2^=0.45, p-value=0.0002). Each circle indicates a bundle, the two hemispheres are presented in different shades. Abbreviations: AF: Arcuate Fasciculus, ATR: Anterior Thalamic Radiation, CC: Cingulum Cingulate, CS: Cortico-Spinal Tract, FcMa: Forceps Major, FcMi: Forceps Minor, IFOF: Inferior Frontal Occipital Fasciculus, ILF: Inferior Longitudinal Fasciculus, MLF: Middle Longitudinal Fasciculus, OR: Optic Radiation, SLF: Superior Longitudinal Fasciculus, UNC: Uncinate Fasciculus, VOF: Ventral Occipital Fasciculus, pAF: Posterior Arcuate Fasciculus, L=left, R=right.

In the dHCP data, in which scan age ranged from 29.29 to 44.7 weeks GA (total period: 15.41 weeks), all bundles showed a positive correlation between infants’ age at measurement and T1w/T2w (Supplementary Table S1 and Supplementary Figure S6) in both the white matter bundles (all r^2^>0.40, all p<5.44e^-32^) and their cortical targets (all r^2^>0.38, all p<7.72e^-30^). In the SVBP data, in which scan age ranged from 39.1 to 48.1 weeks GA (total period: 9 weeks), R1 also increased with age, although the effect was not uniformly significant across bundles (Supplementary Table S2 and Supplementary Figure S7) in this more restricted age range (white matter: all r^2^s between 0.00003 and 0.55, all ps between 0.0001 and 0.94, 19 out of 26 bundles were significant; gray matter: all r^2^s between 0.04 and 0.48, all ps between 0.0005 and 0.37, 20 out of 26 bundles were significant). For both T1w/T2w and R1 slopes were steeper in the white matter than the gray matter (T1w/T2w: t=3.82, p-value=0.0008; R1: t=3.41, p=0.002). Interestingly, in a few bundles, these differences in developmental slopes between tissues lead to a crossing over where initially lower white matter values began to exceed gray matter values during development. For example, in the cortico-spinal tract (CS), we observed a cross-over in T1w/T2w at around 37 weeks GA and in the superior longitudinal fasciculus (SLF) we observed a cross-over in R1 at around 43 weeks GA (Supplementary Figure S6 and S7).

Similar to the mean values reported above, the slopes of T1w/T2w and R1 change in gray and white matter also varied across the bundles. For T1w/T2w, the CS, for instance, stood out with the highest rate of change in both the white matter bundle and at its cortical target. In contrast, the ATR, pAF, and the FcMi exhibit comparatively low rates of change, with the other bundles falling in between. For R1, the CS did not stand out as clearly, but the FcMi and ATR again exhibit comparatively low rates of change, with the other bundles falling in between. Further, again similar to the mean values, bilateral bundles showed similar gray and white matter slopes between the left and right hemisphere in both T1w/T2w and R1. As the slopes of T1w/T2w and R1 are derived from datasets with different scan age ranges, they are not directly comparable. However, even when we matched the scan age ranges (by including only those subjects from the dHCP that are within the same range as the SVBP, N=183), we find that T1w/T2w and R1 slopes are not significantly correlated with each other (Supplementary Figure S8) in either the gray matter (r^2^=0.07, p-value=0.20) or the white matter (r^2^=0.06, p-value=0.24).

To determine if there is a relationship between the slope of white and gray matter development, we correlated T1w/T2w as well as R1 slopes of both tissues across bundles (Fig. 3c,d). Our analyses revealed a positive correlation between the slopes of white matter bundles and their cortical targets in both metrics (T1w/T2w: r^2^=0.49, p<0.0001; R1: r^2^=0.45, p-value=0.0002), suggesting coupled white and gray matter development during early infancy. For both metrics, these observed correlations for the true bundle–target pairings were above the 95% confidence intervals (CI) of chance-level correlations (95% CI of chance-level: r^2^= 0.15 for T1w/T2w, r^2^=0.14 for R1) obtained by 1,000 iterations of correlating shuffled bundle-target pairings, omitting the true pairs (Supplementary Figure S9).

### T1w/T2w coupling relates to behavioral outcomes

As described earlier, we found considerable inter-individual differences in the strengths of T1w/T2w coupling across tissues and as such we aimed to further characterize these inter-individual differences in the large-scale dHCP data. We find that the T1w/T2w coupling is dependent on gestational age at birth (r^2^=0.18, p-value=5.15e^-13^, Fig. 4a) and gestational age at scan (r^2^=0.18, p-value=2.23e^-13^, Fig. 4b), but not on sex (t=-1.88, p-value=0.06, Fig. 4c). We also tested whether these inter-individual differences in T1w/T2w coupling correlate with later behavioral outcomes. To do so, we analyzed a subset of infants from the dHCP data (N=215) that also completed the Bayley-III questionnaire, and hence allowed us to link T1w/T2w measurements taken at birth to behavioral outcomes assessed between 17 and 25 months of age. For the three Bayley-III subscales (motor, language, and cognition) we investigated the relationship between each individual’s age-standardized test scores and their T1w/T2w coupling across tissues. For cognition and language, we found no significant link between the T1w/T2w coupling across tissues and performance (all r^2^<0.007, all ps>0.11, Fig, 4d,e), however, we found that the T1w/T2w coupling across tissues correlates with later-life motor performance (r^2^=0.02, p=0.015, significant with a Bonferroni corrected threshold, Figure 4f). We also tested if T1w/T2w measured in gray matter or in white matter alone is linked to later-life behavioral outcomes (Supplementary Figure S10), but found no significant relationship between T1w/T2w in either tissue and any of the three subscales (all r^2^<0.007, all ps>0.23). Inclusion of demographic covariates (birth age, scan age, or sex) did not significantly improve the linear model relating T1w/T2w coupling and motor performance (scan age: *F*(1,212) = 2.58, *p* = 0.11, birth age: *F*(1,212) = 1.09, *p* = 0.30, sex (*F*(1,212) = 1.18, *p* = 0.28).

**Figure 4:**
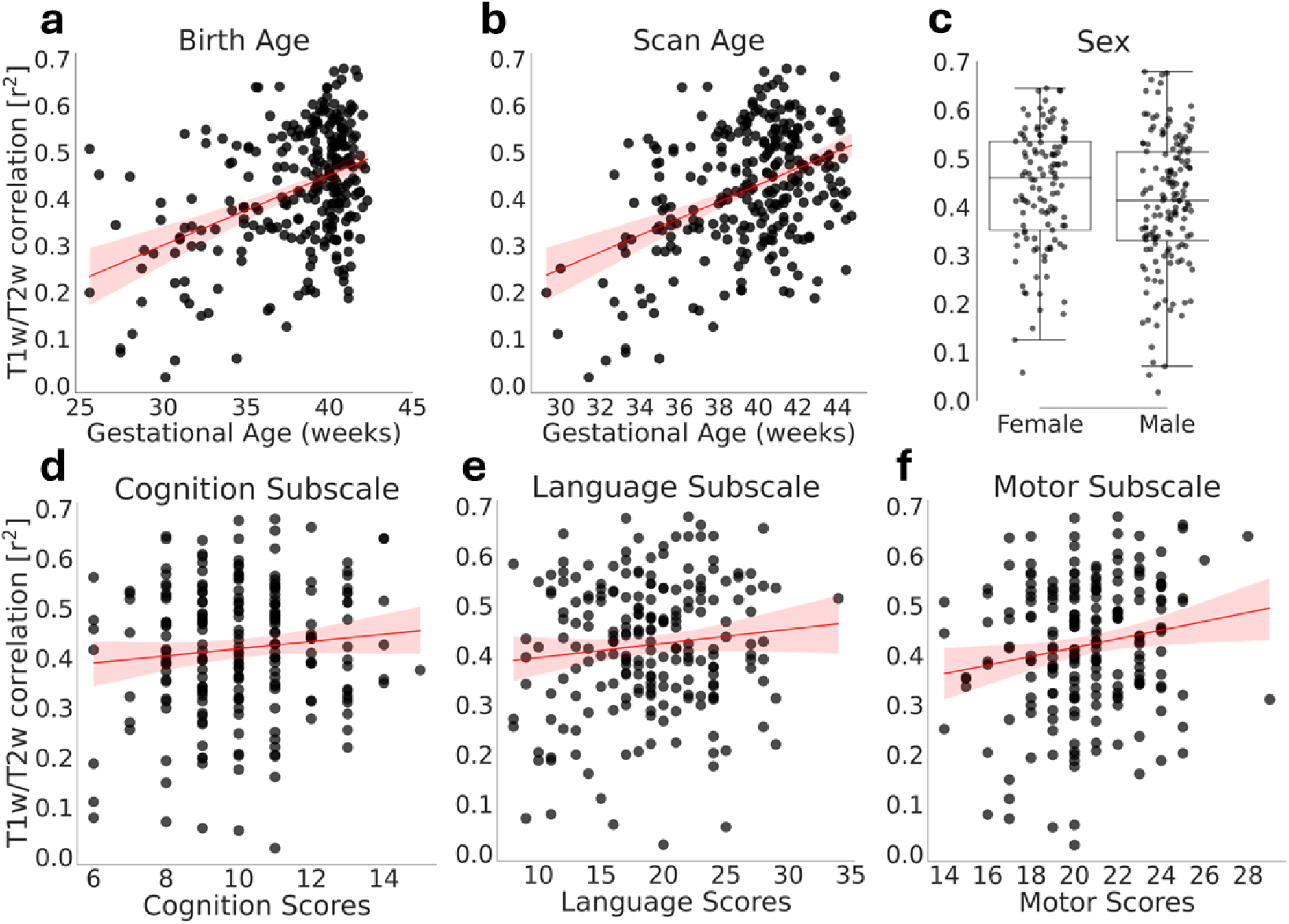
Characterization of inter-individual differences in the T1w/T2w coupling and its relationship to behavior. a-c. Upper row relates T1w/T2w coupling across tissues to gestational age at birth (a, r^2^=0.18, p-value=5.15e^-13^), gestational age at scan (b, r^2^=0.18, p-value=2.23e^-13^), and sex (c, t=-1.88, p-value=0.06). **d-f** Bottom row relates the correlation of T1w/T2w in gray and white matter to the cognition (d; r^2^=0.01, p-value=0.14), language (e; r^2^=0.01, p-value=0.11), and motor (f; r^2^=0.03, p-value=0.02) subscale of the Bayley-III. We found a relationship between the T1w/T2w coupling and motor skills, but not the other subscales. Each dot is a subject, red lines indicate linear regression lines, shaded regions indicate 95% confidence interval, boxes in c indicate the interquartile range (IQR, 25th–75th percentile), the line inside each box denotes the median and the whiskers extend to 1.5×IQR.

### Reduced T1w/T2w coupling in preterm infants

To examine if the coupling of T1w/T2w across white and gray matter is impacted by prematurity, we examined this relationship in a small longitudinal sub-sample from the dHCP that focuses on infants born preterm. This sample contained dMRI and T1w/T2w data from 26 preterm infants (GA at birth: mean ± SD: 32.04 ± 2.97 weeks) scanned once shortly after birth (GA at first scan: mean ± SD: 34.33 ± 1.76 weeks) and once after they reached term-equivalent age (GA at second scan: mean ± SD: 40.70 ± 1.08 weeks), as well as a group of full-term infants with scan ages and sex matched to the preterm infants second scan (GA at scan: mean ± SD: 40.69 ± 1.07 weeks). T1w/T2w of white matter bundles correlated with T1w/T2w of their cortical targets in all three groups: We observed positive correlations in the preterm infants scanned shortly after birth (Fig. 5a; r^2^=0.43, p-value=0.0002), the same preterm infants scanned at term-equivalent age (Fig. 5b; r^2^=0.54, p<0.0001), and full-term infants matched on age and sex to the preterm infants’ second scan (Fig. 5c; r^2^=0.58, p<0.0001). To determine whether the observed differences in correlation coefficients between groups were statistically significant, we employed a non-parametric bootstrap resampling procedure (1,000 iterations) to estimate the sampling distribution of each pairwise difference. The resulting 95% confidence intervals (CIs) were then compared against 0 to assess whether group differences were significant. This analysis revealed weaker correlation between T1w/T2w of white matter bundles and their cortical targets in the preterm infants at their first scan, compared to their full-term peers (Fig. 5d, mean difference in r^2^=-0.10, 0 fell outside 95% CI [-0.101, -0.097]). Similarly, we also found weaker correlations within the preterm infants at their first compared to their second scan (Fig. 5e, mean difference in r^2^ =-0.08, 0 fell outside 95% CI [-0.081, -0.072]). When we compared preterm infants at their second scan to their age-matched full-term peers, we found a higher correlation in the full-term infants (Fig. 5f, mean difference in r^2^=-0.02, 0 fell outside 95% CI [-0.020, -0.016]).

**Figure 5:**
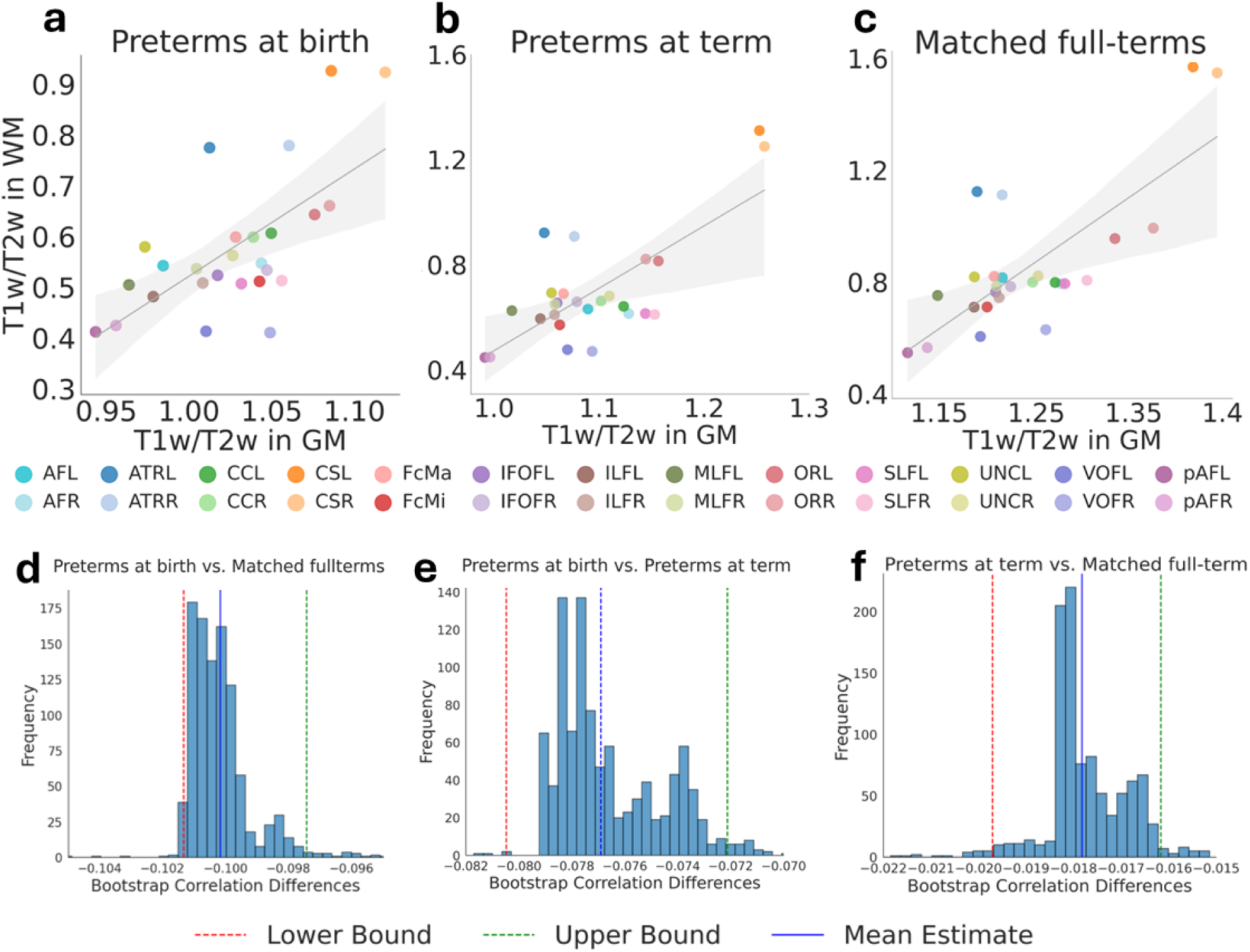
Prematurity impacts the T1w/T2w coupling across tissues. a-c Correlation of T1w/T2w across tissues in three groups: a infants born preterm scanned shortly after birth (r^2^=0.43, p-value=0.0002); **b** infants born preterm scanned at term-equivalent age (i.e. at ≥ 37 weeks gestational age, r^2^=0.54, p<0.0001); **c** age-matched infants born full-term (r^2^ = 0.58; p<0.0001). Each dot is a bundle. **d-f** Bootstrap distributions of correlation differences between groups: **d** preterm infants scanned shortly after birth vs. full-term infants; **e** preterm infants scanned shortly after birth vs. preterm infants scanned at term-equivalent age; **f** preterm infants scanned at term-equivalent age vs. full-term infants. The vertical red and green lines indicate the upper and lower bound of the 95% confidence intervals (CIs), blue line indicates mean. Zero fell outside the 95% CIs in all comparisons. Abbreviations: AF: Arcuate Fasciculus, ATR: Anterior Thalamic Radiation, CC: Cingulum Cingulate, CS: Cortico-Spinal Tract, FcMa: Forceps Major, FcMi: Forceps Minor, IFOF: Inferior Frontal Occipital Fasciculus, ILF: Inferior Longitudinal Fasciculus, MLF: Middle Longitudinal Fasciculus, OR: Optic Radiation, SLF: Superior Longitudinal Fasciculus, UNC: Uncinate Fasciculus, VOF: Ventral Occipital Fasciculus, pAF: Posterior Arcuate Fasciculus, L=left, R=right.

Further comparisons between the three groups revealed a few additional interesting patterns. First, bundles show an overall similar order of development across the groups: the corticospinal tract (CS) exhibits highest T1w/T2w in both white and gray matter in each group, while the posterior arcuate fasciculus (pAF) shows consistently low T1w/T2w, with the other bundles falling in between. Second, across the groups, the cortical targets of the ATR have lower T1w/T2w and the cortical targets of the VOF have higher T1w/T2w than expected based on their white matter T1w/T2w. Third, there are differences in the degree to which bilateral bundles show similar T1w/T2w values in gray matter across the age group: bilateral bundles are more similar at the preterm infants second scan compared to their first scan (group difference in delta of T1w/T2w across hemispheres: t = 3.54, p=0.0002) and compared to full-term infants (t = -2.79, p=0.01), while the preterm infants first scan and the full-term infants were not significantly different (t = -0.65, p=0.52).

## Discussion

In this study, we investigated the association between myelin-sensitive imaging metrics of white matter bundles and their corresponding cortical targets in two complementary infant datasets: A large-scale dataset shared openly by the dHCP that contained dMRI and T1w/T2w data and a quantitative dataset collected locally at Stanford University that contained dMRI and R1 data. Our analyses revealed five key findings: (i) both T1w/T2w and R1 of white matter bundles are positively correlated with their respective cortical targets; (ii) in both T1w/T2w and R1 the rate of change (slope) is correlated between white and gray matter; (iii) there is substantial inter-individual variability in the degree of T1w/T2w and R1 coupling across tissues; (iv) inter-individual variability in T1w/T2w coupling is related to gestational age and correlates with motor performance at 17-25 months of age; (v) the T1w/T2w coupling is lower in preterm compared to full-term born infants, even at term-equivalent age. Overall, these results suggest that white and gray matter myelination proceed in concert during early infancy.

Here we find that T1w/T2w and R1 development is tightly linked between white matter bundles and their corresponding cortical targets. We hypothesize that this observation might be accounted for by the theory of activity-dependent myelination^25,75^: According to this theory, active neurons send trophic signals (e.g., glutamatergic release, ATP signaling) along their axons that promote oligodendrocyte differentiation both within the cortex and along white matter projections, such that increased neuronal firing can increase myelination in both compartments^76–79^ (for review see^80^). Alternative hypotheses for the observed coupling in myelination include: 1) Coordinated genetic regulation: As similar genetic programs direct the proliferation, migration, and differentiation of oligodendrocyte precursor cells (OPCs) in both cortex and white matter^81,82^, these genetic influences might drive the coordinated myelination of white and gray matter. 2) Common metabolic and microenvironmental factors: Both cortical and white matter myelination are energy-demanding processes that depend on a supply of lipids and other metabolic substrates^18^. As such, the local availability of these resources, coupled with support from glial cells like astrocytes could concurrently influence myelination in both tissues^83,84^.

Our findings relate to prior work linking other gray and white matter properties in different developmental populations: For example, during childhood and adolescence (ages 5-23), the mean diffusivity of white matter bundles is closely linked to mean diffusivity of their cortical targets^85^. White matter properties are also linked to cortical thickness and cortical expansion from childhood to adulthood^86–90^. Further, in adults, white matter connectivity, in combination with cortical microstructure, can be used to predict the functional organization of cortex^91^ and training-induced changes in functional properties of cortex are also associated with white matter changes^92,93^. Studies linking gray and white matter properties during infancy are limited to date, however research showed that white matter connectivity is more strongly impacted by a cortical region’s cytoarchitecture than by its functional properties across infancy, childhood, and adulthood^94,95^. While not specific to changes in myelin, these studies across developmental populations underscore the importance of conjointly evaluating white and gray matter tissue properties.

By simultaneously evaluating changes in myelin-sensitive imaging metrics in both white and gray matter, we had the opportunity to directly compare growth rates across tissues. In both T1w/T2w and R1 slopes were steeper in the white matter than the gray matter during early infancy. These findings can be corroborated when integrating previous research using R1, as R1 is directly comparable across studies. Across studies, R1 growth rates reported for white matter (approximate R1 increase of 0.026s^-1^/month)^5^, exceed R1 growth rates reported for cortex (approximate R1 increase of 0.015s^-1^/month)^4^ during the first 6 months of life, suggesting that white matter myelin content increases faster during early infancy than cortical myelin content. In the present study, we expanded upon this prior work by directly comparing developmental growth rates across both tissues within the same individuals. Interestingly, we observed a crossing over of myelin-sensitive imaging metrics in several bundles, where values were higher in cortex in younger participants but higher in the white matter in older participants. This observation likely links to the commonly-observed inversion of imaging contrasts between gray and white matter during infancy^96^ and provides an interesting direction for future research that could explore inter-individual differences in the timing of this inversion.

In the current study, we combined large-scale assessments of T1w/T2w with assessments of R1 derived from qMRI and found coupled gray and white matter development in both metrics. While there is no direct histological evidence confirming the link between T1w/T2w and myelin content, R1 has been shown to be directly linked to myelin in post-mortem brain tissue^67^. In fact, most studies aiming to validate T1w/T2w as a marker for myelin have used quantitative MRI metrics such as R1 as a gold-standard for comparison. These studies found that while T1w/T2w shows strong test-retest reliability^70^ and aligns well with quantitative measures in adult cortex^74,97^ and infant white matter^59^, it does not align well with quantitative myelin measures in adult white matter^71,72^. This inconsistency is underscored by a recent histological analysis of the corpus callosum^73^, which raised further doubts about the validity of using T1w/T2w as a marker of adult white matter myelination. Our findings add a new perspective to this by showing that the mean values of T1w/T2w and R1 are correlated in early infancy, while their developmental slopes are not, which may point to a strong but non-linear relationship between the two metrics in early infancy (also see^97^). Nonetheless, further research is necessary to fully evaluate the validity of using T1w/T2w as a marker for myelin across different age groups and brain tissues. This validation is especially crucial considering that large-scale initiatives, such as the dHCP, often acquire T1w and T2w images but not quantitative myelin metrics - inducing a trade-off between sample size and measurement validation.

Interestingly, we find that inter-individual differences in T1w/T2w coupling across tissues correlates with later life motor outcomes. Multiple studies have linked increased myelination of either white or gray matter to motor performance in humans^80,98^ and other species such as mice^77,81,99–102^. Our findings complement this literature by showing that it may not only be the absolute amount of myelin that impacts motor performance, but also the degree of synchrony in myelination across white matter bundles and their corresponding gray matter targets. However, as the observed impact of T1w/T2w coupling on motor performance is comparatively small and the coupling itself is impacted by gestational age, future work is needed to assess the implications of this effect. These future studies could relate T1w/T2w coupling to behavioral assessments conducted concurrently with MRI measurements, as here we are predicting motor performance at 17-25 months of age from neuroimaging data collected at birth, which is inherently ambitious. In addition to motor performance, we also explored the link between inter-individual differences in T1w/T2w coupling and later life language and cognitive skills, but found no significant relationship. One possible explanation for our findings being restricted to motor performance may be the time at which the behavioral assessments took place (17-25 months of age), as early infancy is characterized by a particularly rapid attainment of key motor milestones. In fact, most fine and gross motor skills - such as grasping, reaching, and postural control - are typically acquired between 6 and 12 months of age, with movement beginning to serve adaptive functions from 3 to 4 months post-term^103^. Future studies could explore a link between T1w/T2w coupling and language and cognitive abilities later during development.

Our data suggest that the synchrony of gray and white matter myelination may be dependent on brain maturity. First, we find that both gestational age at birth and gestational age at scan are positively related to inter-individual differences in the degree of T1w/T2w coupling across tissues. Further, in a longitudinal sub-sample, we find lower correlation of T1w/T2w across gray and white matter in preterm compared to full-term infants at birth.. Interestingly, even at term-equivalent age, when the age-matched preterm and full-term born infants have had the same amount of time to mature, they still show different degrees of T1w/T2w coupling with coupling being stronger in the full-term born infants. These results are consistent with prior work indicating that preterm birth can alter both the speed and degree of brain myelination, potentially leading to developmental differences when compared to full-term infants^59,104,105^ (for review see^106^).

An important limitation of this study is its reliance on mean values across entire white matter bundles, both cortical target regions, and over all cortical endpoints. By reducing each bundle to a single weighted average value, we are not able to detect any spatial heterogeneity that may exist along its length. Likewise, computing a single weighted average T1w/T2w or R1 value across all cortical endpoints fails to capture localized differences at specific cortical subregions, thereby precluding the detection of fine-grained patterns in cortical myeloarchitecture. This approach may also obscure microstructural variations across different cortical layers, as prior research has shown that T1w/T2w values can vary across cortical depths^22,47,52^. Our analyses thus provide only a birds-eye view of the coupled myelin development across tissues. Future studies employing more granular, regionally, or laminar-resolved T1w/T2w or R1 analyses could provide a more nuanced understanding of this relationship.

Overall, our study leveraged two complementary myelin-sensitive imaging metrics to reveal a tight coupling of myelin development across white and gray matter during early infancy. Recognizing that gray and white matter myelination unfolds in concert compels a rethinking of how we evaluate early life brain development both in research and in clinical settings. Ultimately, acknowledging and further testing the gray-white matter myelination synchrony will refine our understanding of early life brain development and the mechanisms that drive the rapid establishment of myelin across the infant brain.

## Methods

### Participants

For this project, we used anatomical, diffusion-weighted, and T1w/T2w infant data provided by the Developing Human Connectome Project (dHCP) (second data release). Details on data collection parameters are provided directly by the dHCP (http://www.developingconnectome.org/data-release/second-data-release/release-notes/).

The dataset encompassed 490 sessions, acquired from 445 individuals, which included all necessary data for the current analyses (diffusion MRI, T1-weighted, and T2-weighted images). After quality assurance (described below), the dHCP dataset comprised 311 sessions from 273 individuals, including both preterm (gestational age < 37 weeks) and full-term born infants. Of these, 117 were female (Race and ethnicity: 94 White British, 53 White Other, 38 Black/Black British - African, 17 Chinese, 14 Other, 13 Asian/Asian British - Indian, 7 Asian/Asian British - Other, 7 Black/Black British - Caribbean, 6 Any Other Mixed Ethnic Group, 6 White Irish, 6 Unknown, 3 Asian/Asian British - Bangladeshi, 3 Asian/Asian British - Pakistani, 2 White and Black Caribbean, 2 White And Asian, 1 White and Black African, 1 Black/Black British - Other participants). The gestational age at birth ranged from 25.57 to 42.29 weeks (mean ± SD: 38.05 ± 3.67 weeks). Scans were performed between 29.29 and 44.71 weeks post-conceptional age (mean ± SD: 39.53 ± 3.11 weeks), with the time interval between birth and scan ranging from 0 to 16.42 weeks (mean ± SD: 1.48 ± 2.02 weeks). If not explicitly stated otherwise, only the cross-sectional data (the first scan of each individual) was used in the analyses. In addition to this main cross-sectional sample, we also derived two smaller subsamples: The first subsample included cross-sectional data only from those 215 infants (93 females, 55 infants born preterm) that complete the Bayley-III Scales of Infant and Toddler Development, which was collected between 17 and 25 months of age (mean ± SD: 19.15 ± 1.39 months). The second subsample took advantage of the longitudinal data and consisted of two groups. The first group included 26 preterm infants (8 females) who were scanned twice: once shortly after their preterm birth and again at term-equivalent age. For these infants, the mean gestational age at birth was 32.04 ± 2.97 weeks. The first scan occurred at a mean age of 34.33 ± 1.76 weeks, while the second scan was performed at a mean age of 40.70 ± 1.08 weeks. The second group comprised 26 full-term infants matched 1:1 to the preterm group in terms of sex and age at the second scan. This full-term group included 8 females, with a mean gestational age at birth of 40.03 ± 0.82 weeks and a mean scan age of 40.67 ± 1.07 weeks. Study protocols for the dHCP data were approved by the United Kingdom Health Research Authority.

In addition, we also used anatomical, diffusion-weighted and R1 infant data collected locally at Stanford University. We refer to this data as the Stanford VPNL Baby Project (SVBP). Details on data collection parameters are described in^4,5,95^. The data encompassed 27 sessions, acquired from 27 individuals, which included all necessary data for the current analyses (diffusion MRI, R1 maps, and anatomical data). After quality assurance (described below) the data contained 21 infants scanned shortly after birth (Race and ethnicity: 3 Asian, 3 Hispanic, 5 Multiracial, and 10 White participants). Among these participants, 7 were female. The gestational age at birth ranged from 36.5 to 42 weeks (mean ± SD: 39.09 ± 1.63 weeks). Imaging was conducted between 39.10 and 48.10 weeks post-conceptional age (mean ± SD: 43.39 ± 2.53 weeks), corresponding to an interval of 2.3 to 7.1 weeks after birth (mean ± SD: 4.30 ± 1.35 weeks). Study protocols for these data were approved by the Stanford University Internal Review Board on Human Subjects Research.

### Preprocessing

#### dHCP

The anatomical and diffusion-weighted data used in this project underwent several preprocessing steps as part of the dHCP preprocessing pipelines^107–109^ and here we used the preprocessed data. As accurate alignment of dMRI and anatomical data was critical for this project, we applied an additional rigid-body alignment using mrregister (from MRrtrix3^110^) and visually inspected all images for alignment quality. We also used the T1w/T2w maps provided by the dHCP^111^. Information on the generation of these maps are provided directly by the dHCP^112^; very briefly, the T1w image was first rigidly registered to the T2w image and the T1w/T2w was then estimated from the original T2w image and the transformed T1w image (prior to bias correction). Following this, the T1/T2 ratio was projected onto the midthickness surface, using volume-to-surface mapping.

#### SVBP

Preprocessing steps of the SVBP data are described in detail in our prior work^4,5,95^. Briefly, T1- and T2-weighted images for each participant were rigidly aligned and tissue segmentation into gray and white matter was performed with iBEAT V2.0^113^, followed by manual correction in ITKgray^114^. The edited segmentations were then used to reconstruct cortical surfaces with Infant FreeSurfer^115^.

Diffusion MRI data were preprocessed in MRtrix3^110^ (https://github.com/MRtrix3/mrtrix3). Data were denoised using principal component analysis^116^, and susceptibility distortions were corrected with FSL’s topup using a reversed phase-encoded image. Motion and eddy-current artifacts were corrected with FSL’s eddy, which also detected and replaced outlier slices^117^. Bias field correction was applied using ANTs^118^ (https://picsl.upenn.edu/software/ants/). The resulting diffusion data were rigidly registered to the T2-weighted anatomical image, and all alignments were visually inspected.

IR-EPI data were used to estimate R1 (R1 = 1/T1) in each voxel. First, susceptibility-induced distortions were corrected with FSL’s top-up and the IR-EPI acquisition with reverse-phase encoding direction. The distortion corrected images were then used to fit the T1 relaxation signal model using a multi-dimensional Levenberg-Marquardt algorithm^119^. From the T1 estimate, we calculated R1 (R1 = 1/T1) at each voxel. R1 was projected onto the cortical surface using volume-to-surface mapping in FreeSurfer.

### Tractography

Diffusion MRI tractography was conducted similarly for the dHCP and SVBP data while also matching processing steps to our prior work with each of these data sets (dHCP^59^, SVBP^5,95^). Using MRtrix3^110^, first, tissue response functions were computed separately for white matter, gray matter, and CSF using the Dhollander algorithm^120^. We then computed fiber orientation distributions (FODs) with multi-shell multi-tissue constrained spherical deconvolution (CSD)^121^ for the white matter and the CSF. The gray matter was not modeled separately, as the b-values between white and gray matter are not sufficiently different to allow a clean separation of the signals in early infancy. Multi-tissue informed log-domain intensity normalization was performed. A whole brain white matter connectome was then created for each session, and tractography was optimized using the tissue segmentation from anatomical MRI data (anatomically constrained tractography, ACT^122^), which is particularly useful when gray and white matter cannot be separated in the FODs. For each connectome, we used probabilistic fiber tracking with the following parameters: number of streamlines: 2 million (dHCP) or 5 million (SVBP), algorithm: IFOD1 (dHCP) or IFOD 2 (SVBP), step size: 0.2 mm, minimum length: 4 mm, maximum length: 200 mm, FOD amplitude stopping criterion: 0.05, maximum angle: 15 deg. Seeds for tractography were randomly placed within the gray/white matter interface (from anatomical tissue segmentation), which enabled us to ensure that all streamlines reach the gray matter, which was critical for determining each bundle’s cortical terminations.

### Bundle identification with pyBabyAFQ

To assess tissue development across bundles and their cortical targets, it is important to identify the bundles systematically and within the native brain space of each individual infant. In our previous research, we introduced babyAFQ^5,59^, an automated fiber quantification tool specifically designed to identify fiber bundles in the infant brain. BabyAFQ is available both for MATLAB^5^ and for Python^59^ and here we used the Python implementation to identify bundles in the MRtrix-generated whole-brain tractograms from both data sets in a standardized way. As described previously, babyAFQ identifies bundles by using sets of anatomical ROIs that serve as waypoints for each bundle. The way-point ROIs were defined in a newborn template brain^123^ and are converted from this template space to the native brain space of each individual infant. In cases where a streamline crosses the waypoints for multiple bundles, a probabilistic atlas is applied to assess which bundle is the better fit; this probabilistic atlas is also converted from the newborn template space to the brain space of each individual infant. Prior to this work, babyAFQ identified 24 white matter bundles in individual infant’s native brain space (11 in each hemisphere and 2 between-hemispheres): the anterior thalamic radiation (ATR), cortico-spinal tract (CS), forceps major (FcMa), forceps minor (FcMi), arcuate fasciculus (AF), uncinate fasciculus (UNC), superior longitudinal fasciculus (SLF), cingulum cingulate (CC), inferior longitudinal fasciculus (ILF), inferior frontal occipital fasciculus (IFOF), middle longitudinal fasciculus (MLF), ventral occipital fasciculus (VOF) and the posterior arcuate fasciculus (pAF). In the current study, we added the optic radiation (OR) to babyAFQ and ported the pAF and VOF from the matlab version of babyAFQ (Fig. 1). To identify the OR, the central part of the thalamus and the primary visual cortex (V1) served as endpoint ROIs. These ROIs were based on the work of Caffara et al.^124^. An additional way-point ROI was placed in the white matter along the expected OR pathway (between thalamus and V1). All OR ROIs were defined in the same neonate template brain used to identify the other bundles in babyAFQ. For each bundle, we removed streamlines whose trajectories differed by more than four standard deviations from the bundle’s median trajectory, for multiple rounds (see ^125^ for more details). For the large-scale dHCP data cloudknot software^126^ was used to deploy both tractography and bundle identification to the Amazon Web Services (AWS) Batch service.

### Quality assurance

<u>dHCP:</u> For the dHCP data we implemented automated quality assurance steps suitable for a large data set. First, we excluded all sessions that exceeded two standard deviations from the mean with respect to absolute motion and the number of outlier slices replaced by FSL’s eddy as in our prior work^59^. Eight additional sessions were excluded due to conspicuous image artifacts or alignment errors that were noted during data processing. Further, to ensure bundle quality, we excluded all sessions where one or more white matter bundles contained 10 or fewer streamlines, as in our prior work^59^. Overall 167 of 490 sessions were excluded due to low data quality.

<u>SVBP:</u> For the SVBP data we implemented a different set of quality assurance steps, as this smaller data set allowed for manual visual inspections of all data. First, consistent with prior work^4,5,95^, we quantified the number of outlier slices replaced by FSL’s eddy and excluded all sessions with more than 5% outlier slices. No sessions were excluded based on this criteria. Next, we visually inspected all dMRI and R1 data for quality, which led to the removal of six sessions due to conspicuous image artifacts or alignment errors. We also visually inspected all bundles of all individuals to confirm that they adhere to their expected anatomical trajectory. No sessions were excluded based on bundle quality issues. Overall, 6 of the 27 sessions were excluded due to data quality concerns.

### Assessing myelin-sensitive imaging metrics across tissues

After identifying the bundles with pyBabyAFQ, we evaluated T1w/T2w and R1 along their length. For this, each bundle was divided into 100 nodes, and a weighted average of the myelin-sensitive imaging metrics was computed at each node. In this weighted average, the contribution of a particular streamline depends on the Mahalanobis distance of that streamline to the core of the bundle (the median xyz position of that node). That is, streamlines that are closer to the core of the bundle are weighted more heavily than streamlines that are further away from the core. This is customary in the AFQ method and has two important advantages: (i) streamlines closer to the core are more likely to be accurately assigned and less influenced by partial volume effects and therefore provide a cleaner measure of the bundle’s properties, and (ii) the weighting counteracts the effects of bundle volume on the generated bundle profiles. Importantly, in order to ensure that partial voluming between gray and white matter does not contribute to any observed coupling of R1 or T1w/T2w across tissues, the first and last 5 nodes, i.e. the superficial white matter underneath cortex, was excluded from all analyses.

To assess the myelin-sensitive imaging metrics at the cortical targets of each bundle, first we mapped the endpoints of each streamline of each bundle at the gray-white matter interface, using the tckmap function provided by MRtrix3 (Fig. 6). Subsequently, the resulting endpoint density maps were projected onto the cortical surface using FreeSurfer. The same segmentations were employed for both bundle-endpoint identification and cortical-surface generation. For association and inter-hemispheric bundles, which travel from cortex to cortex, this procedure resulted in two cortical areas with endpoints for a given bundle. To distinguish these two cortical areas, in the dHCP data, labels from the draw-EM segmentation were used to define boundaries, so that the origin and termination areas of each bundle could be analysed individually. In the SVBP data, the Desikan-Killiany atlas was used for boundary definition. Notably, we also included three projection bundles (the anterior thalamic radiation, the cortico-spinal tract, and the optic radiation) which only have a single cortical termination area. The cortical T1w/T2w (dHCP) or R1 (SVBP) maps were employed to calculate T1w/T2w or R1 values for every cortical voxel that contains an endpoint density > 0.0001. We then calculated a weighted average across all voxels that contained endpoints for a given bundle, such that voxels with more endpoints also contributed more strongly to the T1w/T2w or R1 values for that bundle’s cortical terminations.

**Figure 6:**
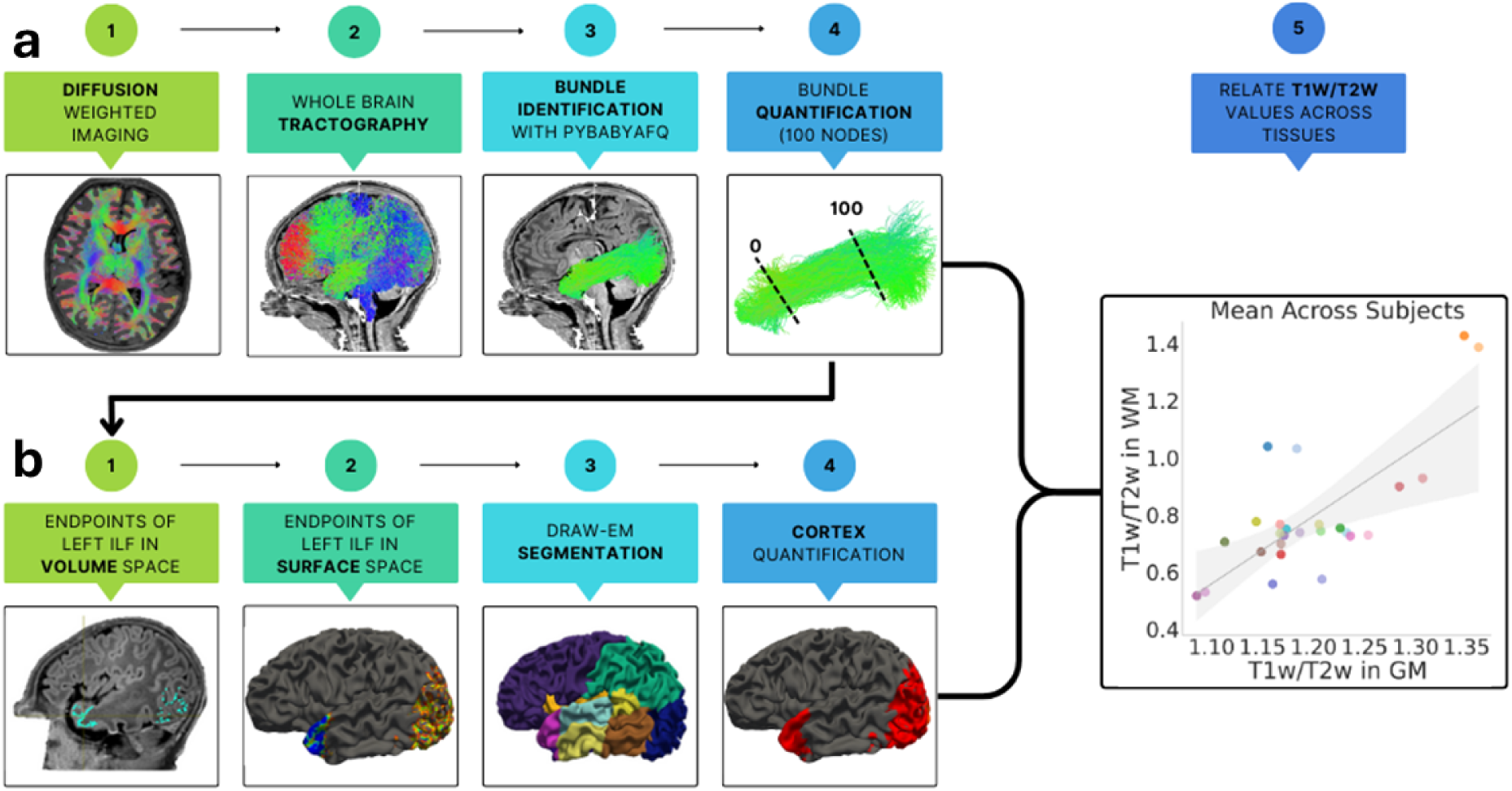
Workflow to obtain white matter (a) and cortical (b) myelin-sensitive imaging metrics. The T1w/T2w of the left inferior longitudinal fasciculus (ILF) is used as an example. White matter T1w/T2w values (top row): A whole brain tractogram is created from dMRI data (steps 1 and 2). Bundle identification is performed with pyBabyAFQ (step 3). Each bundle is divided into 100 equidistant nodes and T1w/T2w is calculated at each node by taking a weighted average of each streamline’s properties at that node (step 4). Gray matter T1w/T2w values (bottom row): After identifying the endpoints in volume (step 1) and surface space (step 2), two cortical endpoint ROIs from the draw-EM segmentation (step 3) serve as anatomical boundaries to separate terminations at the start and end of cortical-to-cortical bundles. At each termination, a weighted average of T1w/T2w is computed based on endpoint density in each voxel (step 4). Finally, T1w/T2w in gray and white matter are related to each other across all bundles (step 5).

### Statistical analyses

We related myelin-sensitive imaging metrics along white matter bundles and their corresponding cortical targets. First, we examined the relationship between mean T1w/T2w (Fig. 2a) and mean R1 (Fig. 2b) in white and gray matter across bundles (for mean values of each bundle see Supplementary Figure S3) using Pearson’s correlation. To determine a chance-level for these correlations, we also computed correlations across tissues in 1,000 iterations of shuffled bundle–target pairings (omitting the true pairings and the homologous bundle in the other hemisphere) and then evaluated if the observed true correlations fell outside the 95% confidence intervals of these random correlations (Supplementary Figure S3). To illustrate inter-individual variability, we also performed subject-wise Pearson correlations and identified two representative cases showing low and high correlation values in each metric (Fig. 2c, d). Additionally, we examined the relationship between T1w/T2w and R1 across bundles in white and gray matter, using Pearson’s correlation (Supplementary Figure S4).

To characterize developmental trajectories, we computed T1w/T2w and R1 slopes for each bundle and their cortical terminations by relating these metrics to gestational age at scan (in weeks) in linear models (models: T1w/T2w ∼ 1 + scan age, R1 ∼ 1 + scan age, Fig. 3a,b, Supplementary Figs. S6, S7 and Supplementary Tables S1 and S2). Paired t-tests were conducted to assess differences in slopes between tissue types. Pearson correlations were employed to test for a relationship between developmental slopes of white matter bundles and their respective cortical targets (Fig. 3c,d). To determine a chance-level for these correlations, we also computed correlations across tissues in 1,000 iterations of shuffled bundle–target pairings (omitting the true pairings and the homologous bundle in the other hemisphere) and then evaluated if the observed true correlations fell outside the 95% confidence intervals of these random correlations (Supplementary Figure S9). We also assessed if the slopes of T1w/T2w and the slopes of R1 are correlated with each other in white and gray matter using Pearson’s correlation (Supplementary Figure S8). For this, in order to ensure that slopes were fit across the same age range, we only included those subjects of the dHCP (N=183) that fell within the (narrower) age range of the SVBP.

As we found large inter-individual differences in the correlation of T1w/T2w across tissues, we aimed to further characterize this inter-individual variability in the large-scale dHCP data. For this we related the strengths of each individual subject’s T1w/T2w coupling, i.e. their correlation in T1w/T2w across tissues, to i) gestational age at birth, ii) gestational age at scan, and iii) sex (Fig. 4a-c). Next we tested if these inter-individual differences have behavioral consequences. For this, we used a subsample of 215 infants from the dHCP which also completed the Bayley Scales of Infant and Toddler Development, Third Edition (Bayley-III^127^), a standardized tool designed to evaluate children from 1 to 42 months of age. Infants whose performance on a given behavioral domain was 2 standard deviations below the mean were excluded to ensure only data from compliant infants was used for the analyses. We examined three behavioral domains (cognition (N=209), language (N=210), and motor (N=205)) by taking the sum of age-standardized scores within each respective Bayley-III subscale and related these behavioral outcomes to the correlation of T1w/T2w across white and gray matter at birth (Fig. 4d-e). In addition, we also tested if T1w/T2w measured in each tissue separately relates to later behavioral outcomes (Supplementary Fig. S10). For each measure, Pearson’s correlations were used to assess associations. Bonferroni correction was applied. Finally, as we found a significant correlation between T1w/T2w coupling across tissues and motor outcomes, we also assess the contribution of demographic variables to this relationship. To this end, we performed a series of nested linear model comparisons. A baseline model predicting standardized Bayley-III motor scores from the T1w/T2w coupling was contrasted against three extended models that each included one additional covariate: scan age, birth age, or sex. All models were fit using ordinary least squares regression implemented in Python (*statsmodels* package). Model comparisons were evaluated using ANOVA-based F-tests, testing whether inclusion of each covariate significantly improves the model fit relative to the baseline model. Statistical significance was set at *p* < 0.05 (two-tailed).

To examine the impact of preterm birth on the correlation of T1w/T2w across tissues, we compared three groups in a small longitudinal sub-sample: (i) preterm infants scanned shortly after birth, (ii) the same preterm infants scanned again at term-aquivalent age, and (iii) full-term infants individually matched to the preterm infants second scan on sex and scan age. For each group, we computed the mean T1w/T2w values in white matter bundles and corresponding cortical targets, and assessed their linear relationship using Pearson’s correlation (Fig. 5a-c). To test if the correlation of T1w/T2w across tissues differs between the groups we employed a nonparametric bootstrapping procedure^128^. For each group, we generated 1,000 bootstrap samples by drawing, with replacement, n = 26 observations and then computed the correlation of T1w/T2w across tissues in each sample. To compare correlations across the three groups, we examined the nonparametric bootstrap distribution of their pairwise differences in r^2^, assessing whether zero lay within or outside the 95% confidence intervals (Fig. 5d-f). When 0 fell outside the confidence intervals, this was interpreted as a significant group difference.

## Data availability

All data required to generate the main figures are provided as a Source Data file with this paper and are also made available in GitHub (https://github.com/EduNeuroLab/WMGMMyelinInfants). Source Data are provided with this paper.

## Code availability

All data were analyzed using open-source software, including MRtrix^110,129^ and pyBabyAFQ, which we shared as a component of pyAFQ (https://tractometry.org/pyAFQ)^125,130,131^. Code that implements the tractography pipeline, example code to perform bundle identification with pyBabyAFQ, and code used to generate the main figures of this manuscript are made available in GitHub (https://github.com/EduNeuroLab/WMGMMyelinInfants).

## Acknowledgements

This work was funded by the European Union (ERC, WRAPPED, 101161197). Views and opinions expressed are however those of the author(s) only and do not necessarily reflect those of the European Union or the European Research Council. Neither the European Union nor the granting authority can be held responsible for them. This work was further supported by the Deutsche Forschungsgemeinschaft (German Research Foundation, DFG) under Germany’s Excellence Strategy (EXC 3066/1 “The Adaptive Mind”, Project No. 533717223) and by the Deutsche Forschungsgemeinschaft (DFG, German Research Foundation) - project number 222641018 - SFB/TRR 135 TP C10, as well as by a LOEWE Professorship awarded by the State of Hesse (LOEWE/4b//519/05/01.002(0016)/120). This research was also supported by National Institutes of Health grants MH121868, MH121867, R01EY033835, and R01EB027585, as well as by National Science Foundation grant 1934292 and by a Stanford Wu Tsai Neurodevelopment grant. Open-source software development was supported by the Chan Zuckerberg Initiative’s Essential Open Source Software for Science program, the Alfred P. Sloan Foundation and the Gordon & Betty Moore Foundation. JK was supported through the NSF Graduate Research Fellowship DGE-2140004. Cloud computing resources were supported by the Amazon Web Services Research Credits Program. Data were provided by the developing Human Connectome Project (KCL-Imperial-Oxford Consortium), funded by the European Research Council under the European Union Seventh Framework Programme (FP/2007-2013), ERC Grant Agreement No. 319456. We sincerely thank the families who generously participated in this study. Data and/or research tools used in the preparation of this manuscript were obtained from the National Institute of Mental Health (NIMH) Data Archive (NDA). NDA is a collaborative informatics system created by the National Institutes of Health to provide a national resource to support and accelerate research in mental health. Dataset identifiers: NIMH Data Archive Collection ID: 3955 or NIMH Data Archive Digital Object Identifier: 10.15154/92vw-g837. This manuscript reflects the views of the authors and may not reflect the opinions or views of the NIH or of the Submitters submitting original data to NDA. We would also like to thank Emily Kubota for assistance in data preprocessing.

## Author contributions

MG, AR and KGS designed the study; CT, XY, ST collected data; SZ, MG, JK, AR, KC, and AO contributed code for data analysis; SZ downloaded, processed and analysed the data; SZ and MG wrote the manuscript. All authors read and provided feedback on the manuscript.

## Competing interests

The authors declare no competing interests.

## Notes

https://github.com/EduNeuroLab/WMGMMyelinInfants

### Competing Interest Statement

The authors have declared no competing interest.

### Summary of Updates

To address reviewers concerns about the validity of T1w/T2w as a myelin proxy, we added analyses from a complementary dataset including diffusion and quantitative MRI, allowing quantification of R1. As R1 is a recognized gold-standard measure of myelin-related changes, its inclusion strengthens our results. Consistent with T1w/T2w, R1 revealed coupled gray and white matter development (Figures 1,2). This addition was conducted in collaboration with colleagues at Stanford University, who are now included as co-authors with approval from all original authors. Including R1 also enabled direct comparison of developmental trajectories between R1 and T1w/T2w in early infancy. Mean values were correlated in both gray and white matter, but their slopes were not, indicating a tight yet non-linear relationship (Supplementary Figures S4, S8). In response to Reviewers comments, we assessed effects of gestational age, scan age, and sex on T1w/T2w coupling and its relation to motor outcomes. Coupling was influenced by age but not sex, and adding these factors did not improve prediction of motor outcomes (Figure 4). To address another question about bundle differences, we added Supplementary Figure S5 and Tables S1,S2, showing mean values and slopes of T1w/T2w and R1 across bundles. Reviewers request for a control analysis led to the inclusion of shuffled bundle target pairs, demonstrating that observed cross-tissue correlations were significantly above chance (Supplementary Figures S3, S9). Finally, we streamlined the manuscript and clarified the Methods section. Supplementary Figure S3 was moved into the main text as Figure 6, and new Supplementary Figures S1,S2 were added to show example bundles across ages and datasets.

