## Supplementary Information for "Cortical and white matter myelination proceed in concert during early infancy"

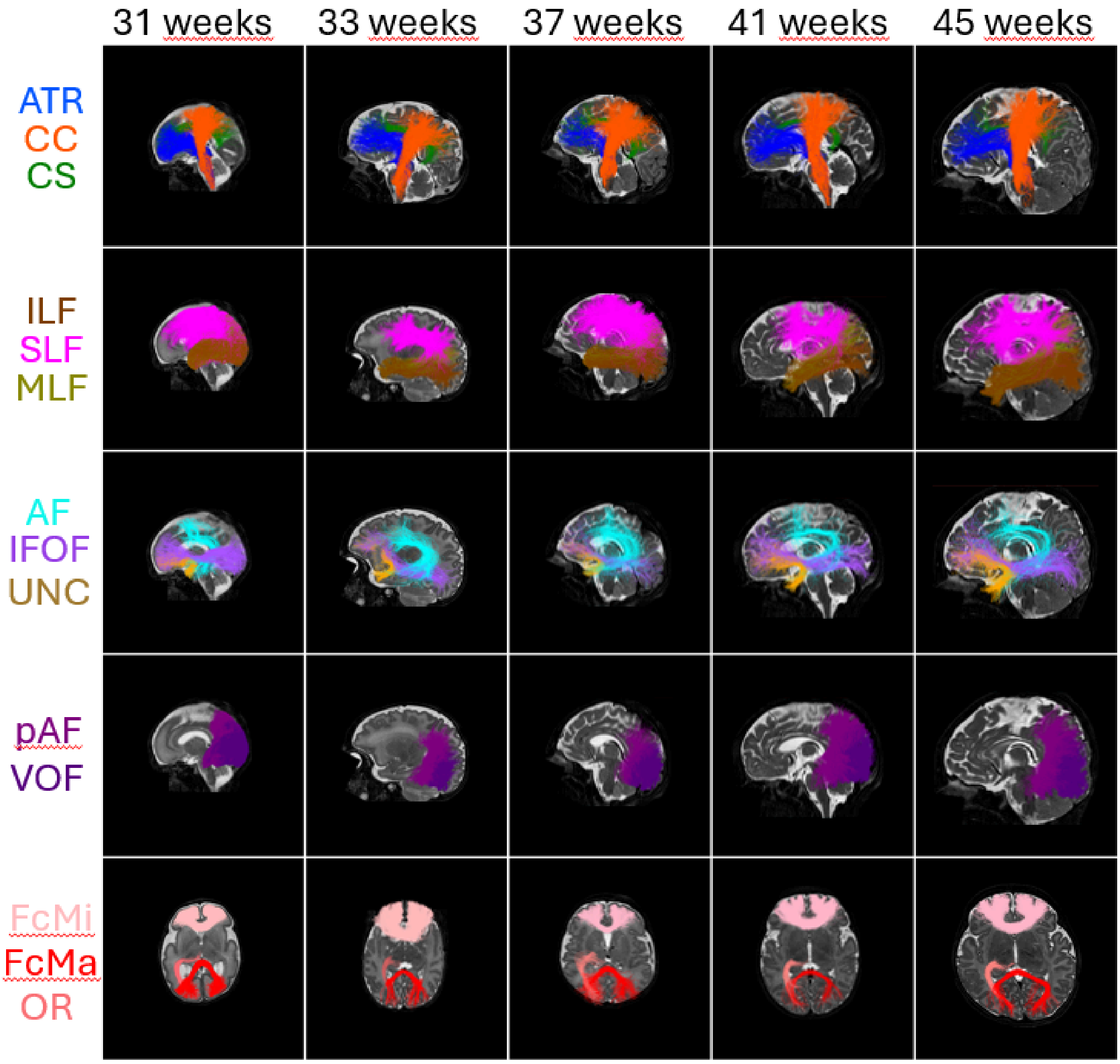


Figure S1: **Examples of bundles identified with pyBabyAFQ in the dHCP dataset.** Bundles are shown in nine randomly chosen individuals from dHCP scanned at gestational ages ranging from 31 weeks (left) to 45 weeks (right). Abbreviations: AF: arcuate fasciculus, ATR: anterior thalamic radiation, CS: cortico-spinal tract, CC: cingulum cingulate, FcMi: forceps minor, FcMa: forceps major, ILF: inferior longitudinal fasciculus, MLF: middle longitudinal fasciculus, IFOF: inferior frontal occipital fasciculus, OR: optic radiation, pAF: posterior arcuate fasciculus, SLF: superior longitudinal fasciculus, UNC: uncinate fasciculus, VOF: ventral occipital fasciculus.

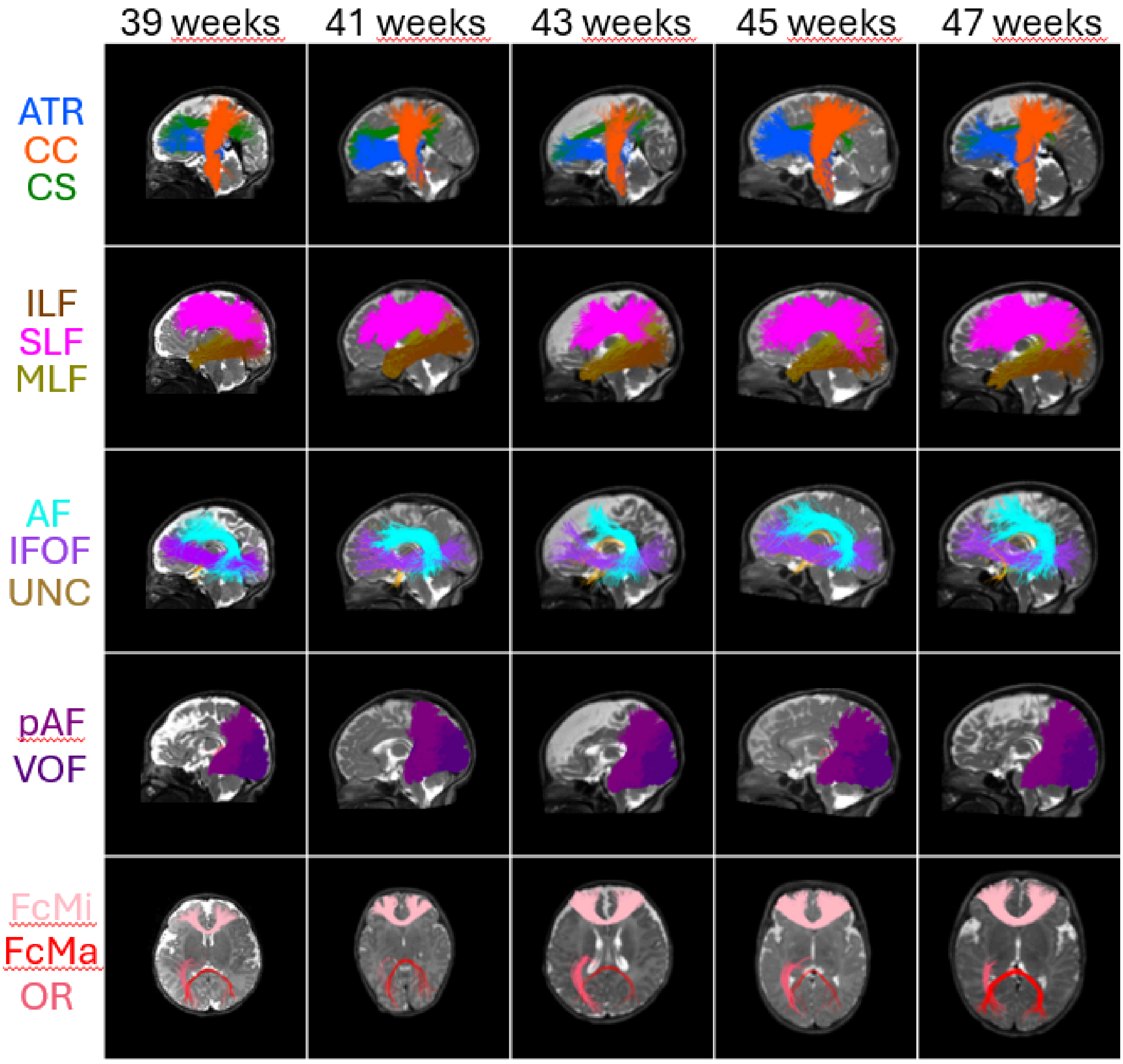


Figures S2: **Examples of bundles identified with pyBabyAFQ in the SVBP dataset.** Bundles are shown in five randomly chosen individuals from SVBP scanned at ages ranging from 39 weeks (left) to 47 weeks (right). Abbreviations: AF: arcuate fasciculus, ATR: anterior thalamic radiation, CS: cortico-spinal tract, CC: cingulum cingulate, FcMi: forceps minor, FcMa: forceps major, ILF: inferior longitudinal fasciculus, MLF: middle longitudinal fasciculus, IFOF: inferior frontal occipital fasciculus, OR: optic radiation, pAF: posterior arcuate fasciculus, SLF: superior longitudinal fasciculus, UNC: uncinate fasciculus, VOF: ventral occipital fasciculus.

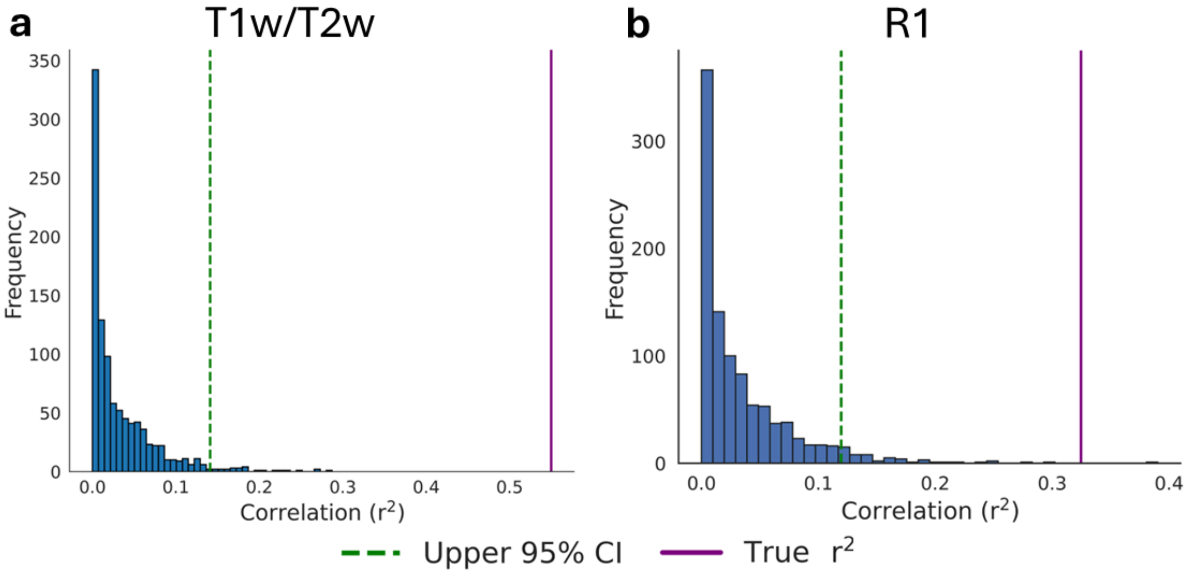


Figure S3: **Results of control analysis.** For this we shuffled bundle–target pairings to estimate chance-level correlations of mean T1w/T2w (a) and R1 (b) values across tissues. The histogram depicts chance correlations observed for 1000 iterations of shuffled bundle–target pairings that omitted the true pairings. The vertical dashed green line indicates the one-sided 95% upper bound of chance-level correlations, and the purple line indicates the observed correlation for the true bundle-target pairings. In both T1w/T2w and R1 the correlations for the true bundle-target pairings are above the 95% one-sided bound.

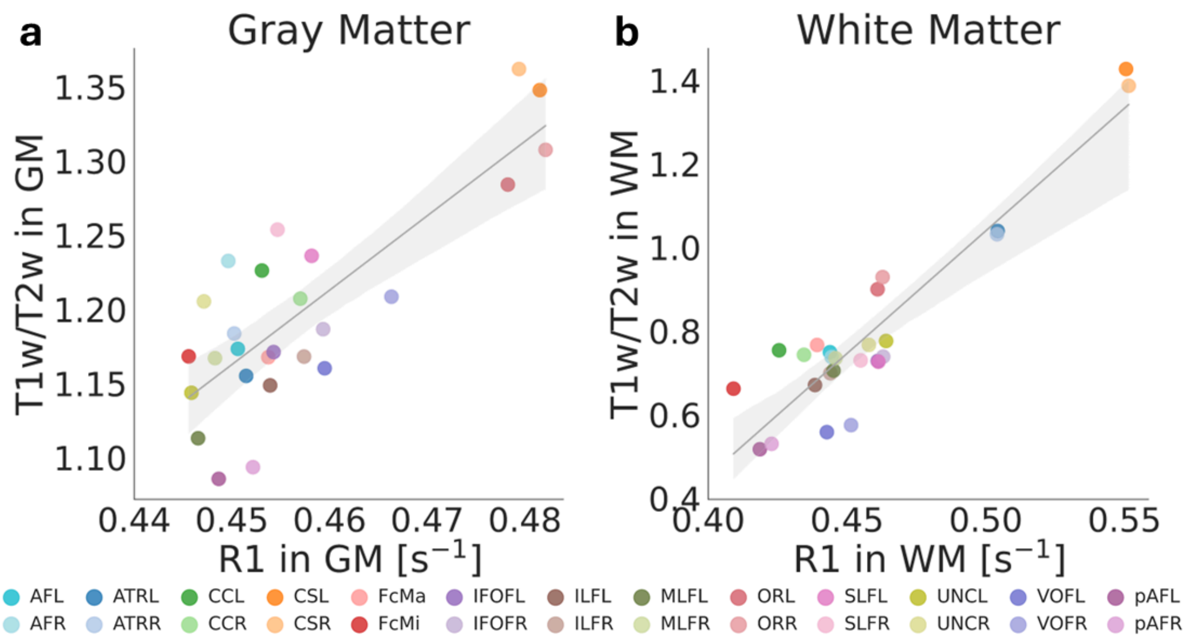


Figure S4: **Relationship between T1w/T2w and R1. a** In gray matter and **b** in white matter. In both tissues the two myelin-sensitive imaging metrics are significantly correlated across bundles (GM: $r^2$=0.66 p-value=5.32e$^{-7}$; WM: $r^2$=0.85, p-value=1.88e$^{-11}$). Values are averaged across subjects, and each dot is a bundle. Abbreviations: GM: gray matter, WM: white matter, AF: Arcuate Fasciculus, ATR: Anterior Thalamic Radiation], CC: Cingulum Cingulate, CS: Cortico-Spinal Tract, FcMa: Forceps Major, FcMi: Forceps Minor, IFO: Inferior Frontal Occipital Fasciculus, ILF: Inferior Longitudinal Fasciculus, MLF: Middle Longitudinal Fasciculus, OR: Optic Radiation, SLF: Superior Longitudinal Fasciculus, UNC: Uncinate Fasciculus, VOF: Ventral Occipital Fasciculus, pAF: Posterior Arcuate Fasciculus, L=left, R=right.

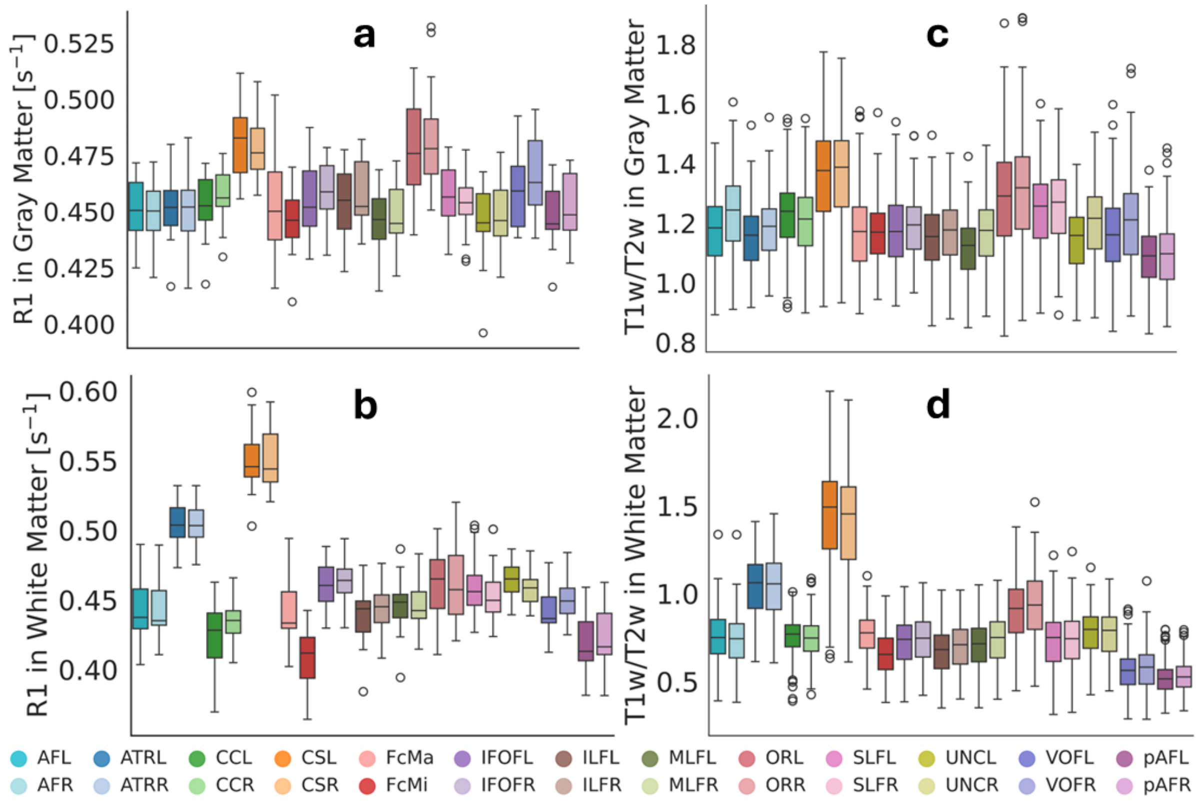


Figure S5: **Boxplots showing differences in T1w/T2w and R1 in white and gray matter across bundles. a,b** R1 in gray matter (a) and white matter (b), respectively. **c,d** T1w/T2w in gray matter (c) and white matter (d) respectively. Each box represents the interquartile range with median (horizontal line), whiskers indicating 1.5 times the interquartile range, and points representing outliers. Colors indicate bundles. Abbreviations: AF: Arcuate Fasciculus, ATR: Anterior Thalamic Radiation], CC: Cingulum Cingulate, CS: Cortico-Spinal Tract, FcMa: Forceps Major, FcMi: Forceps Minor, IFO: Inferior Frontal Occipital Fasciculus, ILF: Inferior Longitudinal Fasciculus, MLF: Middle Longitudinal Fasciculus, OR: Optic Radiation, SLF: Superior Longitudinal Fasciculus, UNC: Uncinate Fasciculus, VOF: Ventral Occipital Fasciculus, pAF: Posterior Arcuate Fasciculus, L=left, R=right.

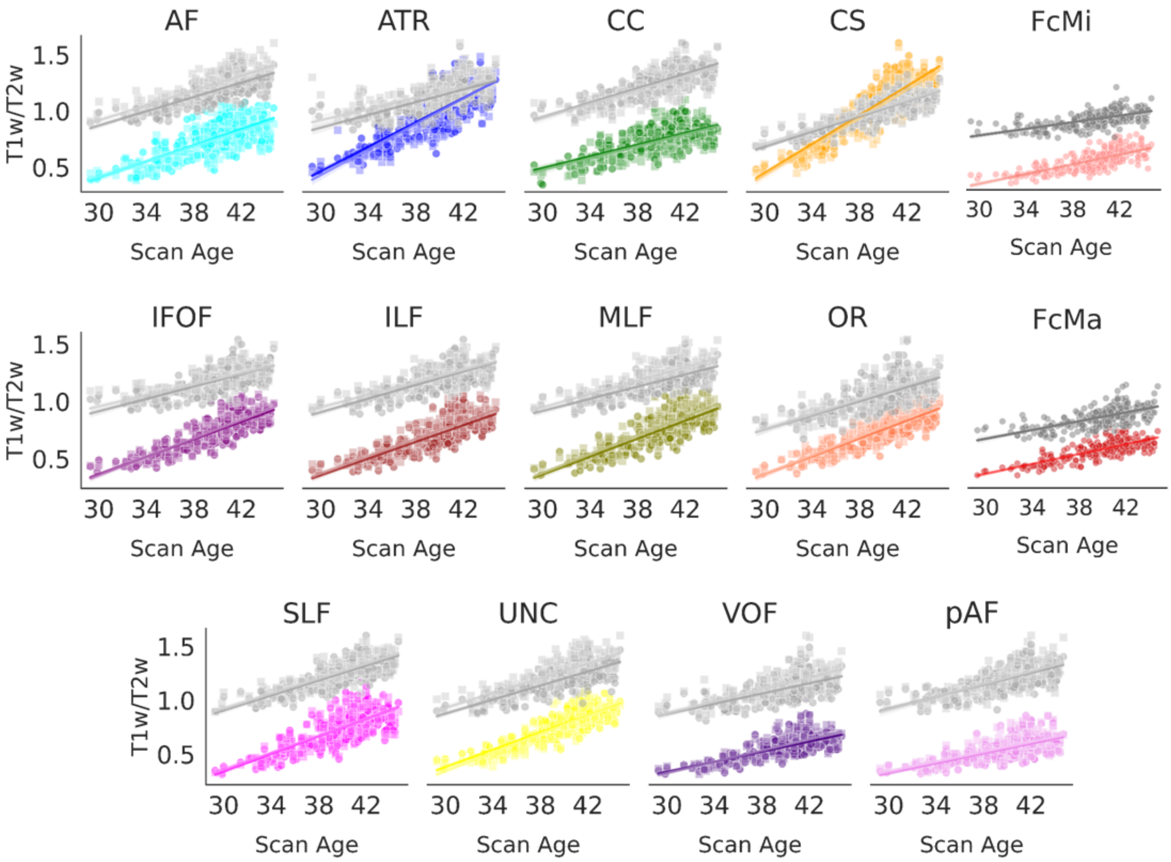


Figure S6: **The rate of change of white and gray matter T1w/T2w across all of the bundles identified with pyBabyAFQ.** Gray markers indicate the correlation for gray matter T1w/T2w, while colored markers represent the correlation for white matter T1w/T2w. Opacity indicates the hemisphere, with lower opacity representing the right hemisphere and higher opacity the left hemisphere. The steepness of the lines indicate slopes of T1w/T2w change with increasing gestational age at measurement in each tissue. Abbreviations: AF: Arcuate Fasciculus, ATR: Anterior Thalamic Radiation], CC: Cingulum Cingulate, CS: Cortico-Spinal Tract, FcMa: Forceps Major, FcMi: Forceps Minor, IFO: Inferior Frontal Occipital Fasciculus, ILF: Inferior Longitudinal Fasciculus, MLF: Middle Longitudinal Fasciculus, OR: Optic Radiation, SLF: Superior Longitudinal Fasciculus, UNC: Uncinate Fasciculus, VOF: Ventral Occipital Fasciculus, pAF: Posterior Arcuate Fasciculus, L=left, R=right.

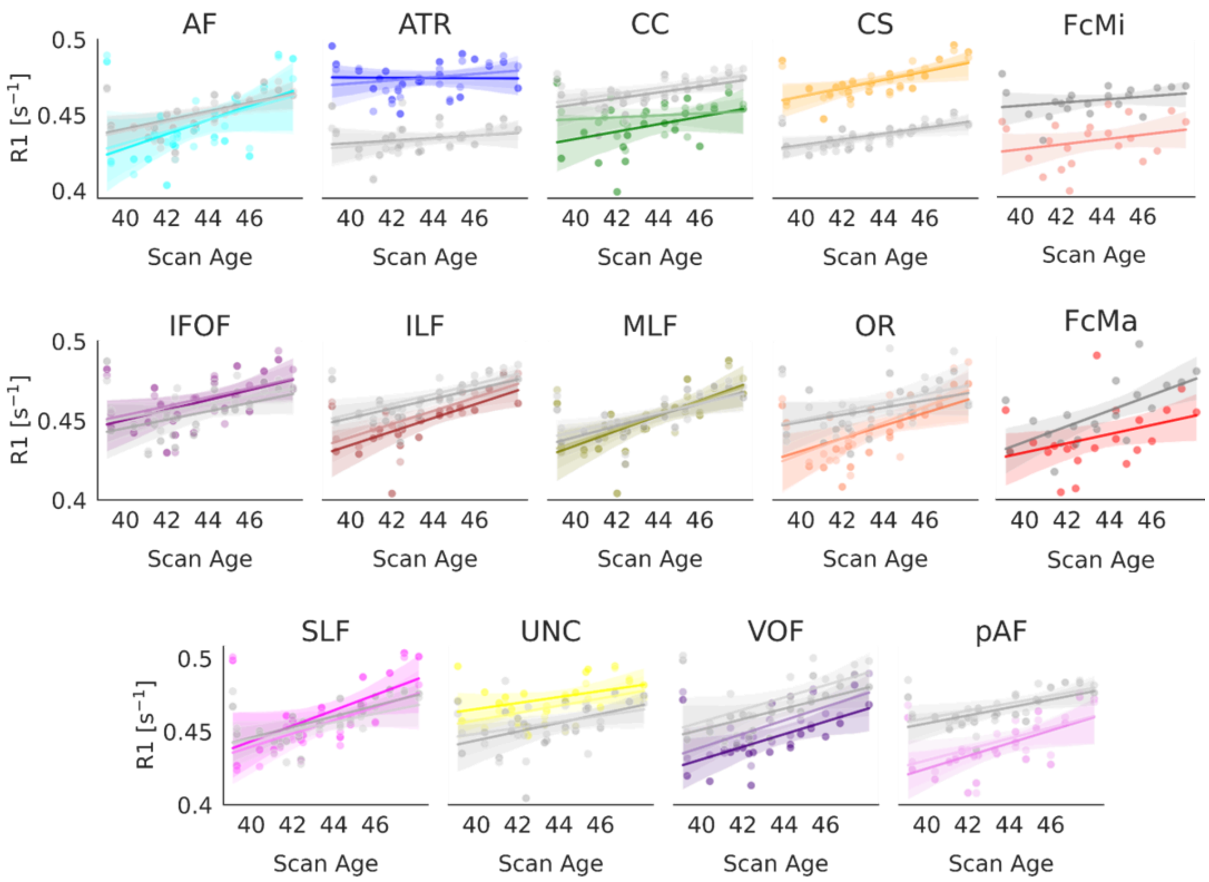


Figure S7: **The rate of change of white and gray matter R1 across all of the bundles identified with pyBabyAFQ.** Gray markers indicate the correlation for gray matter R1, while colored markers represent the correlation for white matter R1. Opacity indicates the hemisphere, with lower opacity representing the right hemisphere and higher opacity the left hemisphere. The steepness of the lines indicate slopes of R1 change with increasing age at measurement in each tissue. Abbreviations: AF: Arcuate Fasciculus, ATR: Anterior Thalamic Radiation], CC: Cingulum Cingulate, CS: Cortico-Spinal Tract, FcMa: Forceps Major, FcMi: Forceps Minor, IFO: Inferior Frontal Occipital Fasciculus, ILF: Inferior Longitudinal Fasciculus, MLF: Middle Longitudinal Fasciculus, OR: Optic Radiation, SLF: Superior Longitudinal Fasciculus, UNC: Uncinate Fasciculus, VOF: Ventral Occipital Fasciculus, pAF: Posterior Arcuate Fasciculus, L=left, R=right.

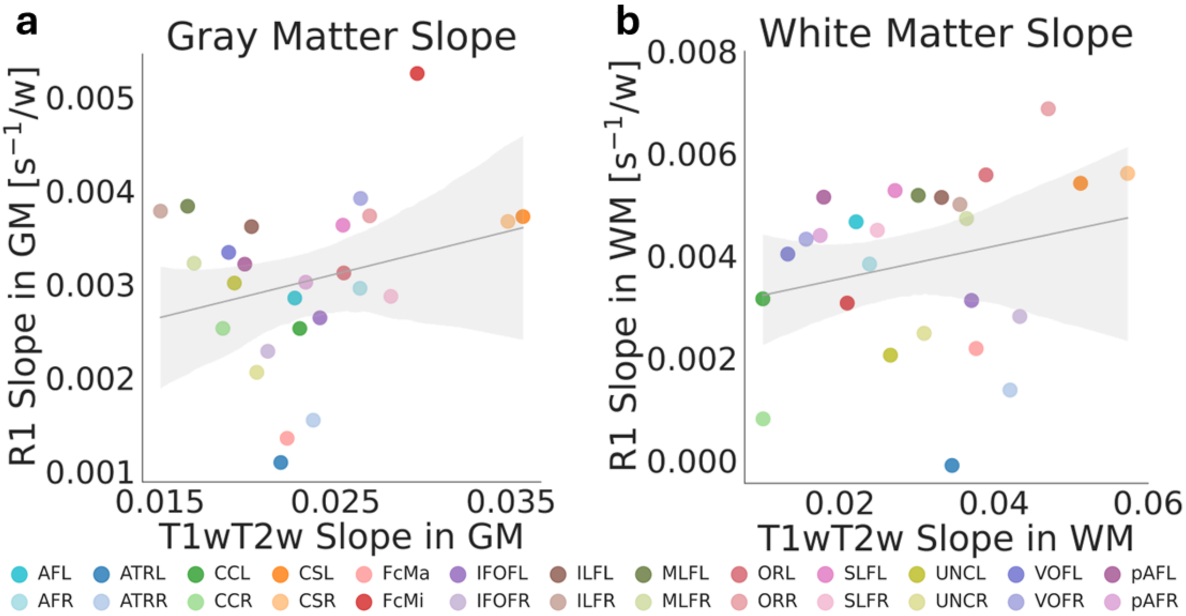


Figure S8: **Relationship between T1w/T2w and R1 developmental slopes in gray matter (a) and white matter (b).** In both tissues the developmental slopes of the two myelin-sensitive imaging metrics are not significantly correlated (GM: $r^2$=0.07, p-value=0.20; WM: $r^2$=0.06, p-value=0.24). The slope indicates the increase in T1w/T2w and R1 values relative to the infants' age at the time of measurement in weeks, each dot is a bundle. Note that we only included those subjects from the dHCP that were within the same age range as the subject of the SVBP (N=183) to ensure that slopes are computed over the same developmental period across metrics. Abbreviations: GM: gray matter, WM: white matter, AF: Arcuate Fasciculus, ATR: Anterior Thalamic Radiation], CC: Cingulum Cingulate, CS: Cortico-Spinal Tract, FcMa: Forceps Major, FcMi: Forceps Minor, IFO: Inferior Frontal Occipital Fasciculus, ILF: Inferior Longitudinal Fasciculus, MLF: Middle Longitudinal Fasciculus, OR: Optic Radiation, SLF: Superior Longitudinal Fasciculus, UNC: Uncinate Fasciculus, VOF: Ventral Occipital Fasciculus, pAF: Posterior Arcuate Fasciculus, L=left, R=right.

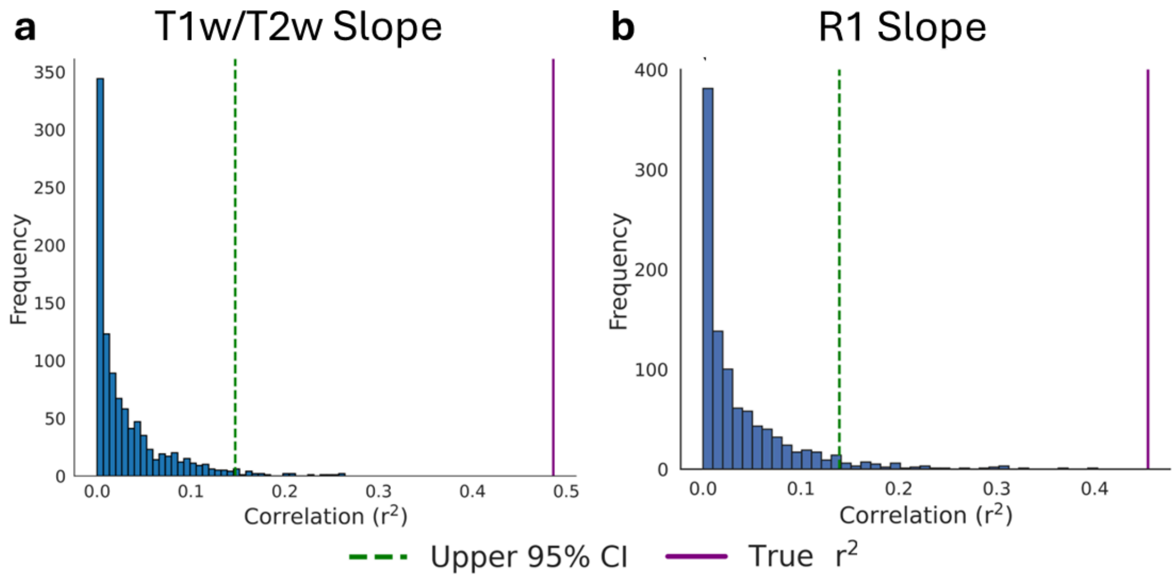


Figure S9: **Results of control analysis for slopes.** For this shuffled bundle–target pairings to estimate chance-level correlations of the slopes of T1w/T2w (a) and R1 (b) development across tissues. The histogram depicts chance correlations observed for 1000 iterations of shuffled bundle–target pairings that omitted the true pairings. The vertical dashed green line indicates the one-sided 95% upper bound of chance-level correlations, and the purple line indicates the observed correlation for the true bundle-target pairings. In both T1w/T2w and R1 the correlations for the true bundle-target pairings are above the 95% one-sided bound.

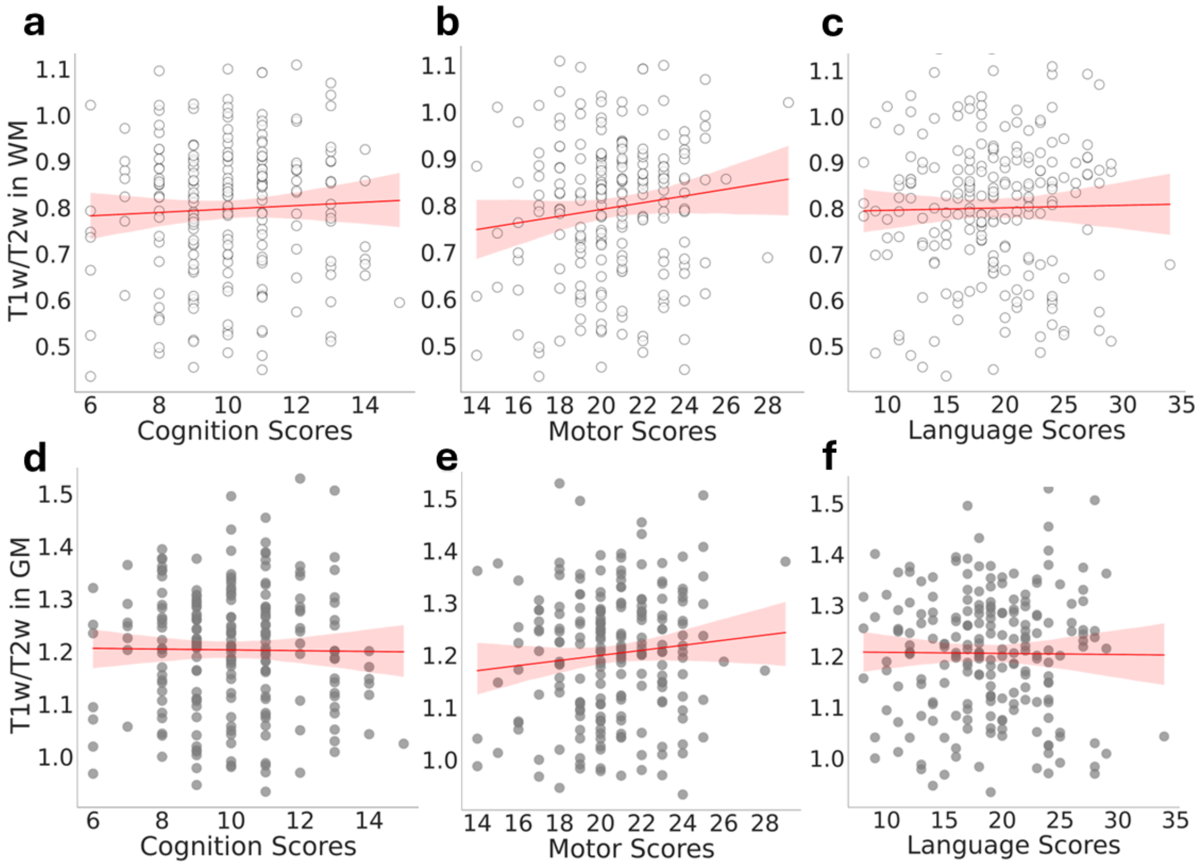


Figure S10: **Relationship between T1w/T2w and cognition, motor, and language performance. a-c** Relationship between T1w/T2w in the white matter and cognition (a; $r^2$=0.002, p-value=0.50), motor (b; $r^2$=0.02, p-value=0.08) and language (c; $r^2$=0.0003, p-value=0.79) subscales of the Bayley-III. **d-f** Relationship between T1w/T2w in the gray matter and cognition (d; $r^2$=-0.0001, p-value=0.87), motor (e; $r^2$=0.01, p-value=0.14), and language (f; $r^2$=0.0008, p-value=0.89) subscales of the Bayley-III.

## T1w/T2w

| Bundle | WM p | WM $r^2$ | WM coeff | GM p | GM $r^2$ | GM coeff |
|---|---|---|---|---|---|---|
| AFL | 3.26E-42 | 0.49601121 | 0.03588858 | 1.01E-62 | 0.64396946 | 0.03183006 |
| AFR | 5.88E-42 | 0.49382503 | 0.0341056 | 4.83E-56 | 0.60128482 | 0.03317838 |
| ATRL | 4.40E-66 | 0.66370581 | 4.48E-02 | 2.18E-38 | 0.46230977 | 0.02336754 |
| ATRR | 7.29E-67 | 0.66813018 | 0.04688158 | 7.72E-30 | 0.3786821 | 0.02065157 |
| CCL | 7.79E-40 | 0.47531305 | 0.02387939 | 2.94E-61 | 0.63502846 | 0.02978607 |
| CCR | 5.44E-32 | 0.40086905 | 0.02437258 | 2.42E-46 | 0.53004997 | 0.02844867 |
| CSL | 1.24E-88 | 0.77062602 | 0.08577072 | 1.77E-74 | 0.7083524 | 0.04492214 |
| CSR | 5.20E-84 | 0.751909 | 0.08429417 | 2.71E-67 | 0.67053776 | 0.04286874 |
| FcMi | 5.04E-48 | 0.54325362 | 0.03047732 | 7.63E-31 | 0.38914321 | 0.02712702 |
| FcMa | 3.30E-56 | 0.6024077 | 0.03014956 | 3.46E-36 | 0.4419104 | 0.02049164 |
| IFOFL | 4.83E-69 | 0.68017531 | 0.03820163 | 2.13E-44 | 0.51432327 | 0.02698286 |
| IFOFR | 3.35E-74 | 0.70697334 | 0.03996037 | 6.77E-42 | 0.49330271 | 0.02417439 |
| ILFL | 2.92E-62 | 0.64118653 | 0.03568442 | 1.29E-50 | 0.56288097 | 0.02914318 |
| ILFR | 1.86E-66 | 0.66583242 | 0.03703395 | 1.13E-47 | 0.54052215 | 0.02678297 |
| MLFL | 2.91E-62 | 0.64119827 | 0.03665657 | 3.90E-50 | 0.55930199 | 0.02484381 |
| MLFR | 1.44E-68 | 0.67759523 | 0.03846351 | 4.64E-49 | 0.55119713 | 0.02753962 |
| ORL | 1.59E-66 | 0.66622128 | 0.04810809 | 5.59E-38 | 0.45857353 | 0.03797427 |
| ORR | 1.15E-65 | 0.6613197 | 0.05121672 | 2.41E-40 | 0.47981063 | 0.04007292 |
| SLFL | 2.46E-52 | 0.57543018 | 0.04031047 | 6.45E-66 | 0.66275742 | 0.03468034 |
| SLFR | 1.30E-51 | 0.57020614 | 0.03956626 | 4.15E-64 | 0.65225524 | 0.03460147 |
| UNCL | 2.67E-71 | 0.69218966 | 0.03503019 | 1.19E-58 | 0.61852777 | 0.02833703 |
| UNCR | 7.06E-80 | 0.73388298 | 0.03747226 | 1.58E-51 | 0.56957773 | 0.02993709 |
| VOFL | 1.12E-45 | 0.52473566 | 0.02609958 | 5.36E-34 | 0.42084628 | 0.02645496 |
| VOFR | 1.13E-48 | 0.54823631 | 0.02909089 | 1.70E-39 | 0.47228797 | 0.03073249 |
| pAFL | 3.70E-39 | 0.46926914 | 0.02012696 | 1.02E-44 | 0.51695343 | 0.0244694 |
| pAFR | 3.09E-38 | 0.46092024 | 0.02004629 | 1.14E-44 | 0.51655634 | 0.02606878 |

Table S1: **Relationship between gestational age and T1w/T2w in each white matter bundle (WM) and their gray matter (GM) targets.** The table summarizes the statistical relationships between scan age (gestational age at measurement in weeks) and T1w/T2w in each bundle. For each bundle, the correlation ($r^2$), the coefficient (slope), and the associated p-value is reported separately for white matter (WM) and gray matter (GM). All bundles show statistically significant correlations (all ps $< 0.05$), as highlighted in green. Abbreviations: GM: gray matter, WM: white matter, AF: Arcuate Fasciculus, ATR: Anterior Thalamic Radiation], CC: Cingulum Cingulate, CS: Cortico-Spinal Tract, FcMa: Forceps Major, FcMi: Forceps Minor, IFO: Inferior Frontal Occipital Fasciculus, ILF: Inferior Longitudinal Fasciculus, MLF: Middle Longitudinal Fasciculus, OR: Optic Radiation, SLF: Superior Longitudinal Fasciculus, UNC: Uncinate Fasciculus, VOF: Ventral Occipital Fasciculus, pAF: Posterior Arcuate Fasciculus, L=left, R=right.

## R1

| Bundle | WM p | WM $r^2$ | WM coeff | GM p | GM $r^2$ | GM coeff |
|---|---|---|---|---|---|---|
| AFL | 0.02197953 | 0.24675132 | 0.00466576 | 0.00649369 | 0.32962908 | 0.00285392 |
| AFR | 0.04100023 | 0.20190967 | 0.00383947 | 0.00934692 | 0.30560913 | 0.00295793 |
| ATRL | 0.94401978 | 0.00026639 | -9.69E-05 | 0.36799227 | 0.0428421 | 0.00109592 |
| ATRR | 0.30297424 | 0.05571395 | 0.0013761 | 0.20932102 | 0.08162375 | 0.00154812 |
| CCL | 0.12518794 | 0.11927374 | 0.00315954 | 0.01885071 | 0.25756295 | 0.00252821 |
| CCR | 0.54563896 | 0.01953842 | 0.00081258 | 0.01501227 | 0.27340477 | 0.00252928 |
| CSL | 0.00349122 | 0.36903122 | 0.00541797 | 0.00162504 | 0.41490455 | 0.00372508 |
| CSR | 0.00060239 | 0.47006738 | 0.00561202 | 0.00269811 | 0.38482106 | 0.00367237 |
| FcMi | 0.24106061 | 0.07156173 | 0.00307622 | 0.25536165 | 0.06750817 | 0.00525478 |
| FcMa | 0.10296567 | 0.13379251 | 0.00218789 | 0.00173945 | 0.41094313 | 0.00135454 |
| IFOFL | 0.0268136 | 0.23261049 | 0.00312842 | 0.04238307 | 0.19948637 | 0.00264276 |
| IFOFR | 0.01881613 | 0.25769162 | 0.00281515 | 0.03487151 | 0.21368809 | 0.00228502 |
| ILFL | 0.00148988 | 0.41992715 | 0.00514167 | 0.00211819 | 0.39934153 | 0.00361781 |
| ILFR | 0.00011579 | 0.55135401 | 0.00500508 | 0.00095596 | 0.44500272 | 0.00378354 |
| MLFL | 0.00132246 | 0.42675957 | 0.00518177 | 0.00054973 | 0.47490963 | 0.00383553 |
| MLFR | 0.00077035 | 0.4568491 | 0.00472878 | 0.00243026 | 0.39113109 | 0.0032256 |
| ORL | 0.00748707 | 0.32031908 | 0.00558123 | 0.10497975 | 0.13235144 | 0.00312207 |
| ORR | 0.00216663 | 0.39799733 | 0.00687402 | 0.05384594 | 0.18190798 | 0.00373258 |
| SLFL | 0.00444538 | 0.35392143 | 0.00527617 | 0.000531 | 0.47673339 | 0.0036329 |
| SLFR | 0.01465996 | 0.27504373 | 0.00449986 | 0.01806566 | 0.26054001 | 0.00287125 |
| UNCL | 0.10039371 | 0.1356746 | 0.00205944 | 0.03977016 | 0.20413202 | 0.00301418 |
| UNCR | 0.01978177 | 0.25417942 | 0.00248636 | 0.12265305 | 0.12079099 | 0.00205944 |
| VOFL | 0.00753326 | 0.31991458 | 0.00403512 | 0.01647233 | 0.26697471 | 0.00334213 |
| VOFR | 0.00179929 | 0.40896507 | 0.00432513 | 0.00824688 | 0.31394034 | 0.00391899 |
| pAFL | 0.00363345 | 0.36655449 | 0.00514678 | 0.00389854 | 0.36216688 | 0.00321604 |
| pAFR | 0.00976442 | 0.30268367 | 0.00439779 | 0.008136 | 0.31483634 | 0.00302265 |

Table S2: **Relationship between gestational age and R1 in each white matter bundle (WM) and their gray matter (GM) targets.** The table summarizes the statistical relationships between scan age (gestational age at measurement in weeks) and R1 in each bundle. For each bundle, the correlation ($r^2$), the coefficient (slope), and the associated p-value is reported separately for white matter (WM) and gray matter (GM). Statistically significant correlations ($p < 0.05$) are highlighted in green. Abbreviations: GM: gray matter, WM: white matter, AF: Arcuate Fasciculus, ATR: Anterior Thalamic Radiation], CC: Cingulum Cingulate, CS: Cortico-Spinal Tract, FcMa: Forceps Major, FcMi: Forceps Minor, IFO: Inferior Frontal Occipital Fasciculus, ILF: Inferior Longitudinal Fasciculus, MLF: Middle Longitudinal Fasciculus, OR: Optic Radiation, SLF: Superior Longitudinal Fasciculus, UNC: Uncinate Fasciculus, VOF: Ventral Occipital Fasciculus, pAF: Posterior Arcuate Fasciculus, L=left, R=right.
